# Targeting cancer-associated fibroblasts for treatment of ER+ breast cancer: A mathematical modeling perspective and optimization of treatment strategies

**DOI:** 10.64898/2026.03.27.714662

**Authors:** Tuğba Akman, Kristian Pietras, Alvaro Köhn-Luque, Ahmet Acar

## Abstract

Cancer-associated fibroblasts (CAFs) are a central component of the tumor microenvironment that facilitate a supportive niche for cancer progression and metastasis. Experimental evidence suggests that CAFs can facilitate estrogen-independent tumor growth, thereby reducing the efficacy of anti-hormonal therapies. Understanding and quantifying the complex interactions between tumor cells, hormonal signalling, and the microenvironment are crucial for designing more effective and individualized treatment strategies. We propose a mathematical framework to explore the influence of CAFs on ER+ breast cancer progression and to evaluate strategies to mitigate their impact. We develop a system of nonlinear ordinary differential equations that substantiates the experimental observations by providing a mechanistic basis for the role of CAFs in regulating estrogen-independent growth dynamics. We then employ optimal control theory to evaluate distinct therapeutic approaches involving monotherapy or combinations of: (i) inhibition of tumor-to-CAF signaling, (ii) inhibition of CAF-to-tumor proliferative signaling, and (iii) endocrine therapy. Taken together, our results demonstrate that CAF-targeted strategies can enhance treatment efficacy across various estrogen dosing regimens. Our study provides new insights into the potential of CAF as a therapeutic target that could help to improve existing approaches for endocrine therapies.

## 1 Introduction

Breast cancer is a heterogeneous disease and remains a leading causes of cancer-related deaths among women worldwide [22]. Estrogen receptor-positive (ER+) breast cancer accounts for approximately 70% of all diagnosed cases [17]. This subtype is defined by the overexpression of estrogen receptor alpha (ER*α*), which is activated upon binding with estradiol (E2) [36]. Consequently, ER*α* signaling serves as the primary driver of tumor cell proliferation and survival [32], which has lead to the development endocrine therapies aimed at disrupting this pathway [41]. However, despite the initial success of these interventions, intrinsic or acquired resistance remains a major therapeutic challenge in a significant number of cases.

The tumor microenvironment (TME) plays a pivotal role in modulating cancer progression and mediating therapeutic resistance [51]. Within TME, cancer-associated fibroblasts (CAFs) have emerged as critical drivers of tumor growth [23, 39, 40]. These stromal cells maintain a dynamic TME through bidirectional signalling with cancer cells [38]. Specifically, CAFs remodel the extracellular matrix, secrete a variety of growth factors, and modulate immune responses [5]. Notably, CAFs have been implicated in reprogramming estrogen receptor signalling in luminal breast cancers [46]. Experimental evidence suggests that CAFs can facilitate estrogen-independent tumor growth, thereby undermining the effectiveness of endocrine therapies and enabling continued proliferation in hormone-deprived environments [42].

Mathematical modelling of breast cancer treatment allows to investigate therapeutic outcomes. For example, Enderling et al. developed a partial differential equation model for tumor growth and invasion to investigate the effect of different radiation treatment modalities [14]. Model simulations suggested that both conventional external beam radiotherapy and targeted intraoperative radiotherapy following breast conserving surgery can successfully eliminate tumor cells that may have escaped from treatment. Schmiester et al. developed a system of ordinary differential equations to predict the combined action of a CDK4 */*6 inhibitor plus endocrine treatment in Luminal B patients using the baseline expression of six genes from a tumor biopsy [45]. Lai et al. simulated personalized chemotherapy treatment using a multi-scale model informed by multi-modal patient data and shed light on individual treatment outcomes [25, 26]. Wu et al. derived a mathematical model to identify patient-specific treatment schedules in triple negative breast cancer patients, using patient-specific MRI data [54]. Benzekry et al. calibrated a mathematical model for neoadjuvant treatment response of primary tumor and metastasis by using a mouse model [3]. Their study reveals that anti-tumor effects of neoadjuvant receptor tyrosine kinase inhibitor treatment can differ between the primary tumor and metastases in the preoperative setting.

The optimization of cancer treatment scheduling, formulated as an optimal control problem (OCP), have received considerable attention [44, 47, 15, 10, 21, 11]. For example, optimal combination treatment with trastuzumab and doxorubicin was designed for HER2+ breast cancer by Lima et al. [31]. Akman et al. developed a mathematical model for ER+ breast cancer using mouse data and investigated optimal anti-hormonal treatment under diet differences [1]. Their results suggest that high-fat diet and obesity influence clinical outcomes. Wang and Schattler designed an OCP in case of tumor heterogeneity and they showed that the optimal protocols could be designed with a period of lower dose drug administration but a full dose therapy segment [52]. Wu et al. developed a personalized model for breast cancer by integrating MRI data to predict the efficacy and toxicity of neoadjuvant therapy via doxorubicin based on computational fluid dynamics [53]. As an intracellular protein modelling for MCF7 cells, Heldring et al. proposed a mechanistic model [20]. They underlined the importance of further research on the pharmacodynamics of endocrine disrupting chemicals. Li and Thirumalai developed a mechanistic model for HER2+ and HER2− breast cancer by taking cellular heterogeneity into consideration [30]. Their study revealed that not only the drug dose, but also the treatment period should be adjusted for better treatment efficacy. Xiong et al. explored the combined treatment of mRNA-based cancer vaccine and anti-CTLA-4 antibody as an OCP [55]. It was shown that there is a positive correlation between the doses of anti-CTLA-4 antibody and the mRNA-based cancer vaccine.

Despite the prevalence of mathematical modeling in breast cancer and treatment optimization, models that explicitly incorporate Cancer-Associated Fibroblasts (CAFs) remain scarce. Heidary et al. proposed an agent-based model that accounts for the dual contradictory role of fibroblasts in tumor progression [19]. Similarly, Lee and Kim explore the phenotypic diversity of CAFs, specifically their opposing effects on tumor growth and immune response [28], noting that distinguishing between these characteristics is key for the success of targeted treatments.

Quantitative understanding of the complex interactions between tumor cells, hormonal signalling, and stromal cells is crucial for designing more effective and individualized treatment modalities. In this study, we propose a mathematical framework exploring the effect of CAFs on ER+ breast cancer progression to investigate treatment strategies mitigating their impact. We first construct a baseline model consisting of nonlinear ordinary differential equations (ODEs) that describe the temporal evolution of tumor volume, ER*α* concentration, and estrogen-bound ER complexes. The model captures key biological processes including logistic tumor growth, transcription and degradation of ER*α*, and estrogen-receptor binding kinetics. This core model was then extended to include the dynamics of CAF populations, incorporating the feedback loop on the proliferative effects on tumor growth and modulatory effects on ER signalling. A mice experiment comparing different levels of E2 supply, conducted by Reid et al. [42] is the main inspiration for the present work. Using experimental data from ER+ breast cancer mouse models subjected to varying levels of estrogen exposure [42], we calibrated the model parameters to reflect individual tumor responses. Notably, we examined treatment outcomes under high, medium, and low doses of estrogen, allowing us to capture a range of physiological and therapeutic conditions. Our model successfully reproduced observed tumor growth dynamics and revealed how variations in estrogen availability and CAF activity jointly influence the progression of the disease. Then, we implemented three different treatment strategies as an optimization problem to reduce the tumor size during treatment. To do this, we focused on inhibition of proliferative effect of tumor cells on CAFs, inhibition of proliferative effect of CAFs on tumor cells and inhibition of E2-ER binding. We compared constant and optimal application of therapy when mono or combination treatment was applied. Taken together, our results demonstrate that incorporating CAF-targeted strategies significantly enhances treatment efficacy across various estrogen dosing regimens. The presented model describes how CAFs are not merely passive bystanders but active modulators of treatment resistance, and hence, their inhibition can substantially reduce tumor growth, even under hormone-depleted conditions. The findings highlight the potential role of CAFs as an intervention strategy and demonstrate that quantitative modelling can bridge the gap between experimental evidence and clinical translation.

## 2 Material and Methods

### 2.1 Data collection

Reid et al. investigated the impact of CAFs on the therapeutic outcomes of breast cancer [42]. Here, we utilize data from their work involving the MCF7 cancer cell line, a well-characterized ER*α*+ luminal model derived from metastatic breast adenocarcinoma. In their experimental setup, MCF7 cells were orthotopically transplanted into NSG mice. Tumor growth was supported by varying levels of exogenous estrogen via 60-day slow release pellets that contained 0.5 mg, 0.1 mg, or 0.025 mg 17*β*-estradiol. The study observed distinct dose-dependent responses. At high dose (0.5 mg), tumor growth occurred regardless of CAF co-injection. At low dose (0.025 mg), MCF7 cells alone failed to establish tumors; whereas co-transplantation with CAF resulted in substantial tumor development. In a different condition, tumors were initially induced with 0.1 mg E2 pellets followed by estrogen withdrawal. Only the tumors co-transplanted with CAFs persisted and continued to grow, confirming that CAFs facilitate estrogen-independent growth in vivo. Experimental groups consisted of five mice each. Tumor volume was measured at seventeen different time points for low dose of E2, twenty-one time points for medium E2 dose and sixteen time points for high E2 dose [42, Fig. 4D]. Additionally, we use the ER*α*-intensity data reported in the study [42, Fig. 4A].

### 2.2 Model

We develop an ODE model that provides a simplified mechanistic description of the interplay between ER+ breast tumor cells, ER*α*, E2ER complexes in the absence or presence of CAFs.

#### 2.2.1 Model without CAFs

First, we build a model for the temporal dynamics of tumor volume *T* := *T*(*t*), ER*α* concentration *ER* := *ER*(*t*) and estrogen-bound ER complexes *E*2*ER* := *E*2*ER*(*t*) in the tumor tissue at time *t*. The model is based on the following assumptions:

- Tumor volume follows logistic growth dynamics [2].
- The E2ER complex mediates cellular proliferation [37].
- While ER*α* translation and degradation are biologically regulated by the ubiquitin-proteasome system, signaling pathway crosstalk and mRNA stability [49, 16], we represent these processes using effective constant rates to maintain model parsimony, consistent with previous literature [20].
- Binding and unbinding of ER*α* and E2 is assumed to follow first-order kinetics with constant rate coefficients [43].
- The metabolic clearance or degradation of E2ER complex occurs at a constant rate [43].
- Initial baseline concentrations of ER*α* and E2ER are assumed to be uniform across all experimental subjects.
- We neglect degradation of E2 and assume the concentration is constant for a given time.

Consequently, we develop the following system of ODEs:

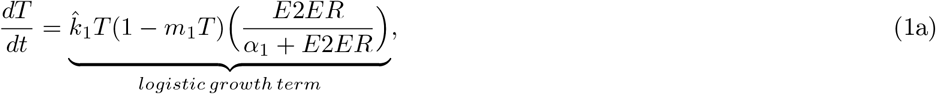

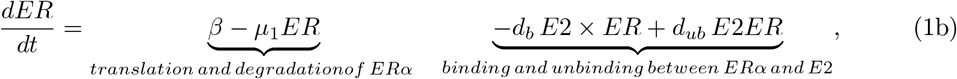

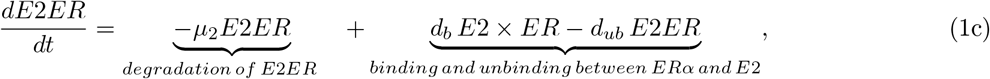

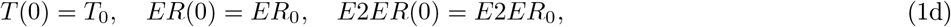

where

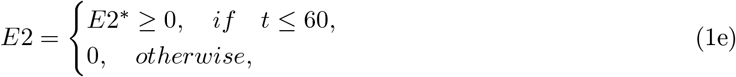

with the non-negative initial conditions *T*_0_, *ER*_0_ and *E*2*ER*_0_. A flow diagram depicting the interactions between model variables *T, ER* and *E*2*ER* is presented in Fig. 1.

**Figure 1:**
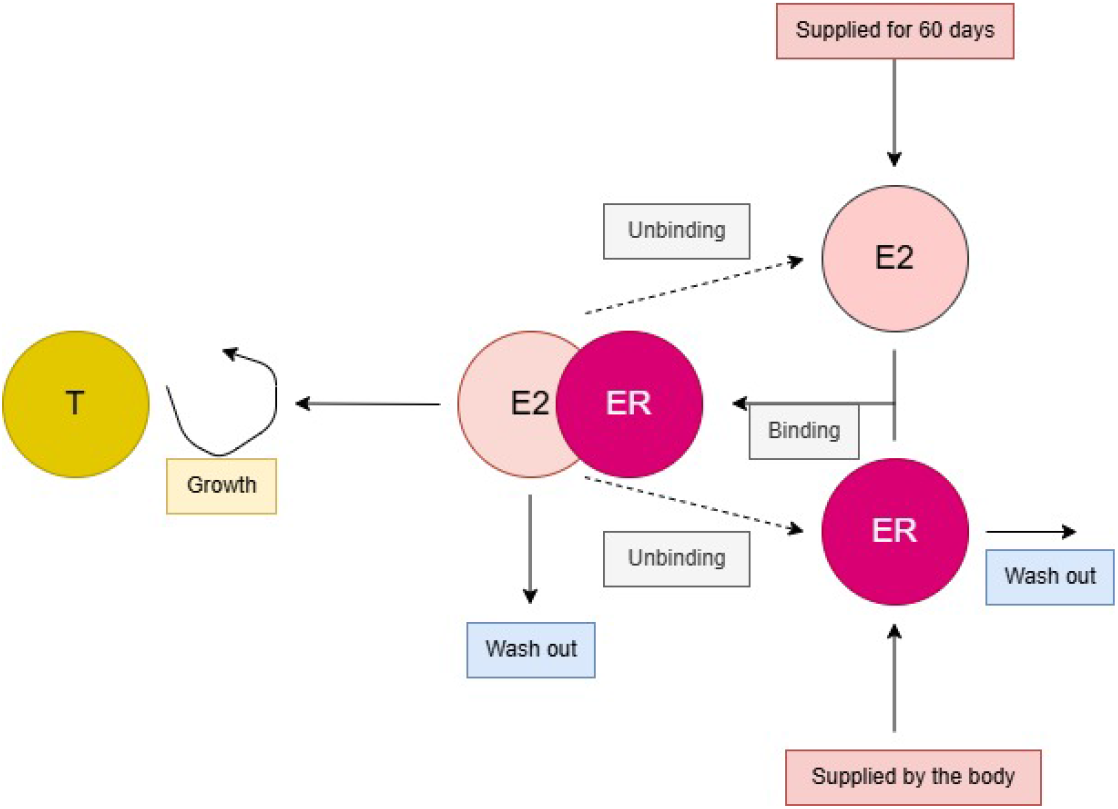
Flow diagram for Model (1).

The parameters 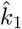, *m*_1_, *α*_1_, *β, µ*_1_, *d*_*b*_, *d*_*ub*_ and *µ*_2_ are all non-negative. Eq. (1a) models the logistic tumor growth, where the growth rate is assumed to follow Michaelis-Menten kinetics depending on the concentration of E2ER complex, 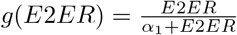. Parameter 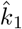 is the maximum growth rate for high E2ER level and *α*_1_ is the E2ER level at which the growth rate is half-maximum. Parameter *m*_1_ is the inverse carrying capacity of the tumor. Eq. (1b) represents the change in ER*α* activity due to its translation and degradation at the rates *β* and *µ*_1_, respectively, and binding and unbinding between ER*α* and E2 occur at the rates of *d*_*b*_ and *d*_*ub*_, respectively. The last equation (1c) accounts for E2ER complex due to binding and unbinding of ER*α* and E2ER. Additionally, E2ER degrades at a rate of *µ*_2_.

#### 2.2.2 Model with CAFs

We extend model (1) by including CAFs under some additional assumptions:

#### Assumptions

- Breast CAFs have an active role in breast cancer proliferation [4].
- CAFs follow logistic growth.
- Growth of CAFs is triggered by tumor cells [48].
- Translation of ER*α* is inhibited by CAFs [42].

As a consequence, Eq. (1) is extended with the CAFs (*CAF* := *CAF*(*t*)) in the tumor tissue at time *t* as follows:

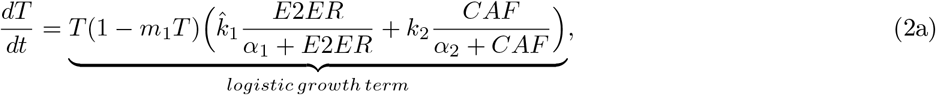

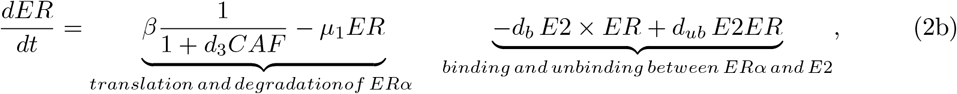

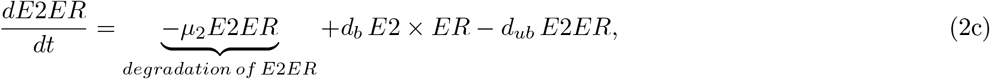

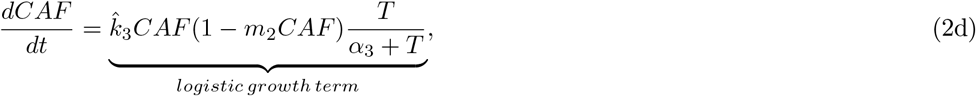

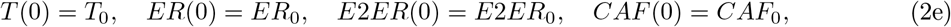

where

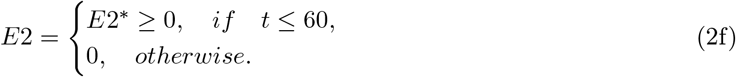

with the non-negative initial conditions *T*_0_, *ER*_0_, *E*2*ER*_0_ and *CAF*_0_.

Eq. (2a) represents the logistic tumor growth and this growth is triggered not only by E2ER complex but also CAFs. Parameters 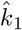 and 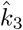 are proliferation rates depending on E2ER complex and CAFs. The growth rates are expressed as 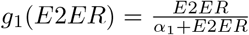 and 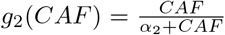. Parameter *α*_2_ is the volume of CAFs at which the growth rate is half-maximum. Eq. (2b) expresses translation of ER*α* inhibited by CAFs. Eq.(2d) describes the logistic growth of CAFs triggered by tumor cells. A flow diagram explaining the interactions between model variables *T, ER, E*2*ER* and *CAF* is presented in Fig. 2.

**Figure 2:**
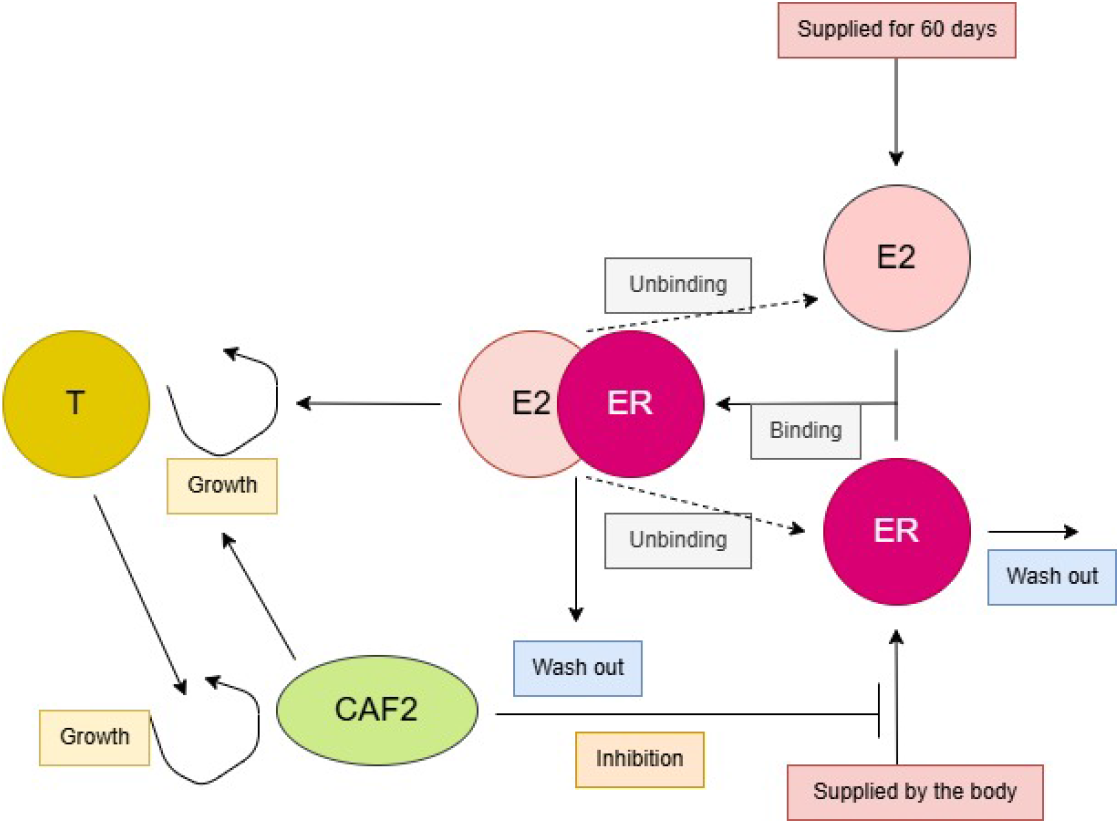
Flow diagram for Model (2).

#### 2.2.3 Structural identifiability analysis

Identifiability analysis has been discussed in many studies (for example [13, 8]). Here, we provide a summary based on these works. There are two types of identifiability that can be investigated: structural and practical identifiability. The former is independent of the data and concerns the structure of the mathematical model, while the latter considers the observations used for model fitting. Some mathematical models suffer from structural nonidentifiability, meaning that a unique parametrization of the model using the available observations cannot be achieved.

We investigate the structural identifiability of model (2) by expressing it as

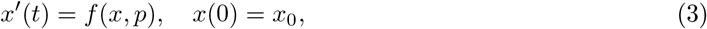

where *x* and *p* denote the states and parameters of model (2), respectively. The observations we have are tumor volume and ER*α* activity, as explained in Sec. 2.1. We can write the observations as [8]

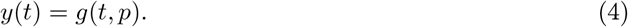

The model given by Equation (3) is structurally identifiable if the vector *p* can be uniquely determined from the observations *y*(*t*) in Equation (4), assuming the observations are unlimited [24]. Otherwise, the model is considered unidentifiable.

Hence, the structural identifiability of the model parameters can be defined as follows [8]:

##### Definition 2.1.

*Let p and* 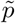 *be distinct model parameter values, and let g*(*t, p*) *be the observations*.

1. *If* 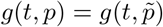 *implies* 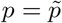, *then we conclude that the model is structurally identifiable from noise-free and continuous observations*.
2. *If* 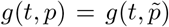 *implies* 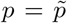 *for any p within an open neighborhood of* 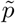 *in the parameter space, then we conclude that the model is locally structurally identifiable*.

It is known from the mice experiment that the human MCF7 cells (0.5 × 10^6^) were implanted either alone or together with human CAFs cells (1.5 × 10^6^) in a 1 : 3 ratio in 100 *µ*l PBS in the 4th mammary fat pad [42]. Therefore, we assume the ratio of the parameters 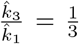 in model (2). This ratio is also needed to overcome structural nonidentifiability in the model (2). From now on, we consider the modified model below:

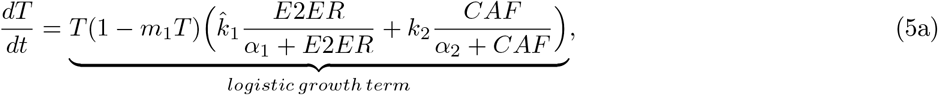

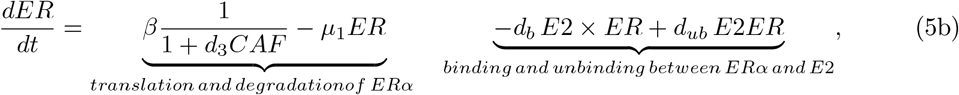

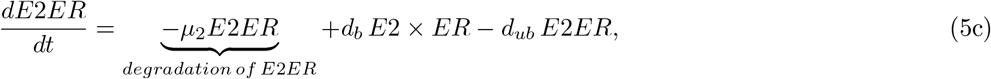

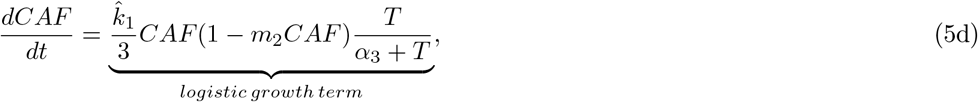

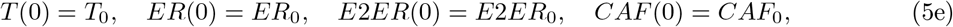

where

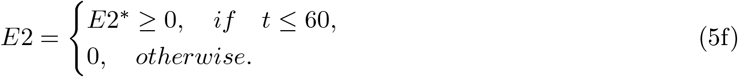

with the non-negative initial conditions *T*_0_, *ER*_0_, *E*2*ER*_0_ and *CAF*_0_.

Structural identifiability can be analyzed using various methods, such as the Lie symmetries [35], generating power series approach [27], and differential algebra approach [34]. In this current work, GenSSI 2.0 is a software toolbox is used [12]. The generating series approach are integrated with identifiability tableaus in the method [6]. Using Lie derivatives of the ODE model, a system of equations is generated. Based on the solvability properties of the system, information about global and local structural identifiability as well as non-identifiability are revealed [7]. The software finds that both Eq. (1) and Eq. (5) are globally structurally identifiable. We present the output of the code in the Appendix.

### 2.3 Parameter estimation

The experimental data described in Sect. 2.1 are used to inform our model (5). We do it by fixing some model parameters using literature values and doing inference on the rest. Inference for model (1) is performed for three different E2 conditions at once by using the data of the mice experiment without CAFs and using Monolix 2024R1 [33]. Random effects of the parameters 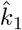, *α*_1_, *β* and *d*_*b*_ are taken into account, while fixed effect associated with the initial values of ER and E2ER are considered. Using average inferred parameters into model (5), we do a second inference using the mice experiment with CAFs for all E2 conditions at once to determine the values of the parameters *k*_2_, *α*_2_, *d*_3_ and *α*_3_ by taking the random effect into account. Next, mean parameter values are used in model (5) to investigate treatment strategies. Parameter values are listed in Table 1.

**Table 1:** Values of the model parameters.

| Parameter | Description | Value | Unit | Source |
| --- | --- | --- | --- | --- |
| $m_1$ | Inverse carrying capacity of tumor cells | 1/1000 | $\text{mm}^{-3}$ | Fixed from the data [42] |
| $m_2$ | Inverse carrying capacity of CAFs | 1/1000 | $\text{mm}^{-3}$ | Assumed |
| $\mu_1$ | Degradation rate of ERα | 0.0042 | $\text{day}^{-1}$ | Fixed [18] |
| $\mu_2$ | Degradation rate of E2ER | 0.0125 | $\text{day}^{-1}$ | Fixed [18] |
| $d_{ub}$ | Unbinding rate between E2 and ERα | 0.0417 | $\text{day}^{-1}$ | Fixed [18] |
| $\alpha_1$ | E2ER threshold for tumor cells to proliferate | 4.45e-5 | - | Calibrated |
| $\alpha_2$ | CAF threshold for tumor cells to proliferate | 21.84 | $\text{mm}^3$ | Calibrated |
| $\alpha_3$ | tumor cells threshold for CAFs to proliferate | 16.28 | $\text{mm}^3$ | Calibrated |
| $\beta$ | Translation rate of ERα | 6.4e-3 | $\text{day}^{-1}$ | Calibrated |
| $d_b$ | Binding rate between E2 and ERα | 1.07e-2 | $\text{mg}^{-1} \text{day}^{-1}$ | Calibrated |
| $d_3$ | Inhibition on ERα by CAFs | 92.75 | $\text{mm}^{-3}$ | Calibrated |
| $\hat{k}_1$ | Growth rate of CAFs | 4.19e-2 | $\text{day}^{-1}$ | Calibrated |
| $\hat{k}_2$ | Growth rate of tumor cells due to CAFs | 0.21 | $\text{day}^{-1}$ | Calibrated |
| E2 | 17β-Estradiol | 0.5, 0.1, 0.025 | mg | [42] |
| $T_0$ | Initial tumor volume | 0.5 | $\text{mm}^3$ | Fixed from the experiment [42] |
| ER$_0$ | Initial ER level | 1.3e-4 | - | Calibrated |
| E2ER$_0$ | Initial estrogen bound estrogen receptor | 3.7e-11 | - | Calibrated |
| CAF$_0$ | Initial CAF volume | 1.5 | $\text{mm}^3$ | Fixed from the experiment [42] |
| $t_f$ | Final time | 160 | days | Assumed based on the experiment [42] |
| $t_{\text{treatment}}$ | Time to start treatment | 45 | days | Fixed from the experiment [42] |
| $u_a$ | Lower bound for control function | 0 | - | - |
| $u_b$ | Upper bound for control function | 0.99 | - | - |

In addition, we present boxplots of the model parameters with respect to three E2 levels in the Appendix (see Fig. 14 and Fig. 15). We observe that the values of *β* and *d*_*b*_ are the same for all mice, while the values of 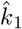 and *α*_1_ vary depending on the E2 level among all mice. We use the average values of the parameters 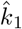, *α*_1_, *β* and *d*_*b*_, since our goal was to construct a general mechanistic model with CAFs. When we display boxplots for the rest of the parameters in the model (5). We note that parameters *k*_2_, *α*_2_, *d*_3_ and *α*_3_ vary depending on E2 supply. We consider average values of the model parameters to assess the contribution of optimal treatment over constant treatment and the effect of the 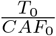 ratio. In addition, temporal evolution of tumor size for each mouse, population fit, prediction bound and the experimental data were found for the models (1) and (5) in the Appendix A (see Fig. 16-23).

### 2.4 Treatment modelling

The next step is to include treatment in model (5). We propose two CAF-targeted treatments via the terms *ũ*_1_ (Type I) and *ũ*_2_ (Type II). The former one targets the growth of CAFs and inhibits it, whereas the latter one targets the tumor growth triggered by CAFs and inhibits it. We inhibit the binding rate of E2 and ER via the term *ũ*_3_ (Type III). For these purposes, we modify model (5) for constant treatment via the constants *ũ*_1_, *ũ*_2_ and *ũ*_3_ and show in Eq. (6). The cases *ũ*_*i*_ = 0 for *i* = 1, 2, 3 correspond to no treatment and their non-negative values model treatment with the inhibition of the corresponding mechanism in Fig. 3.

**Figure 3:**
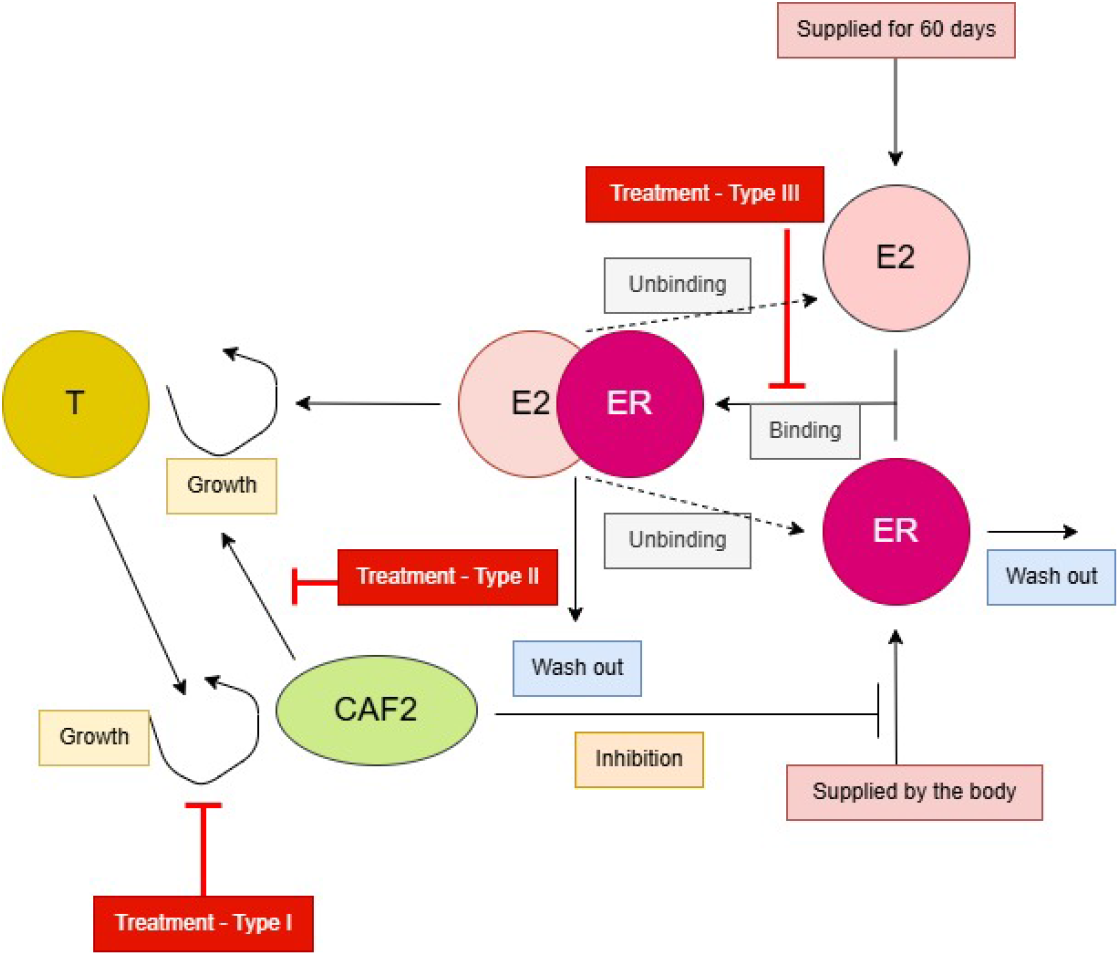
Flow diagram for Model (6).

Then, we obtain the following model for treatment:

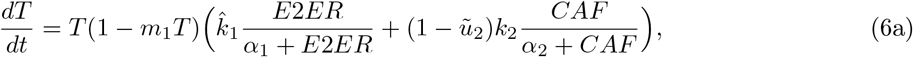

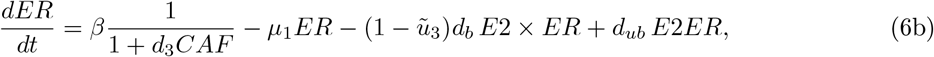

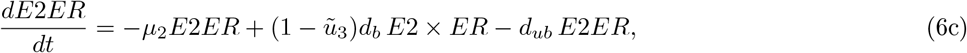

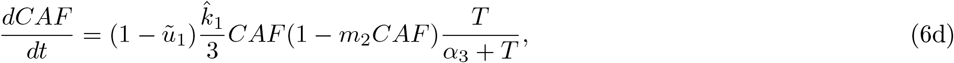

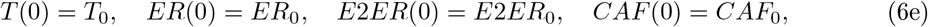

where

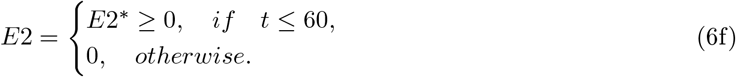

In total, we investigate five different treatment strategies, namely Type I, II, III, combination of Type I and III, combination of Type II and III. Then, we construct an OCP to explore optimal treatment scheduling.

### 2.5 Optimal control formulation

We construct an OCP to minimize the total number of tumor cells together with the side effects of treatment over a pre-specified time interval [*t*_0_, *t*_*treatment*_] with 0 ≤ *t*_0_ *< t*_*treatment*_ ≤ *t*_*f*_. We model the effects of the treatments via time-dependent control functions *u*_1_(*t*), *u*_2_(*t*) and *u*_3_(*t*) without defining the drug dose as a separate model variable. We modify Eq. (6) by replacing the constants *ũ*_1_, *ũ*_2_ and *ũ*_3_ by time-dependent control functions *u*_1_(*t*), *u*_2_(*t*) and *u*_3_(*t*) to construct the optimal control problem

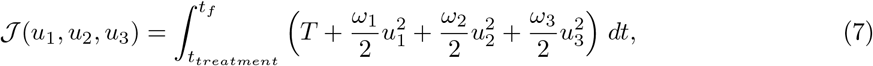

subject to

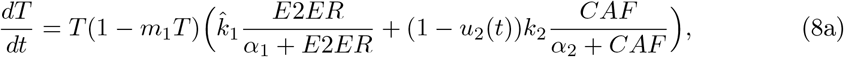

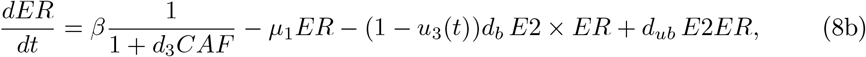

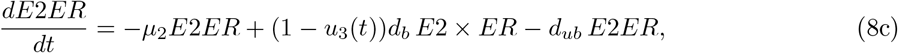

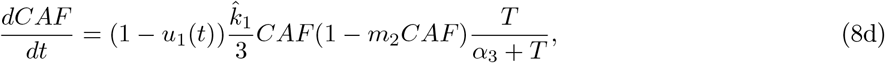

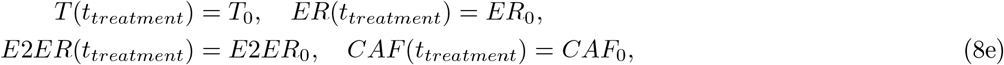

where

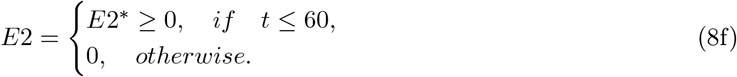

with

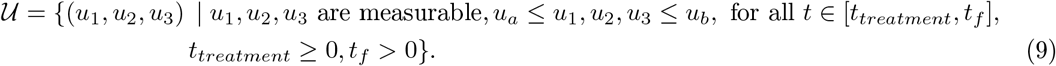

Our aim is to find an optimal control triple 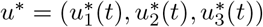 such that

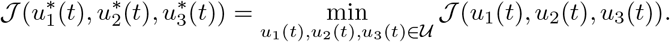

Here, the parameters *ω*_1_, *ω*_2_ and *ω*_3_ in Eq. (7) can be set to balance the size of the different terms. We state the optimality system associated with the OCP (7)-(9).

#### Theorem 2.1.

*Given an optimal control* 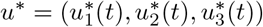 *and a solution* (*T* ^∗^, *ER*^∗^, *E*2*ER*^∗^, *CAF* ^∗^) *to the model* (8) *with non-negative initial conditions that minimizes* (7) *over* 𝒰, *there exist adjoint variables λ*_*i*_ := *λ*_*i*_(*t*) *for i* = 1, …, 4 *such that*

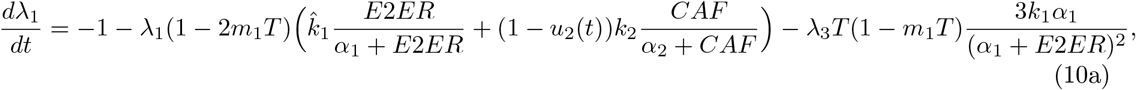

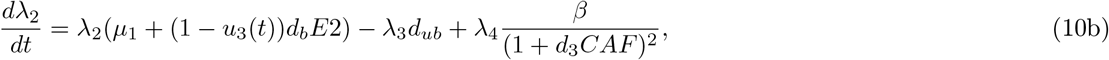

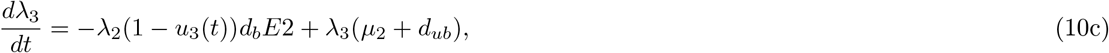

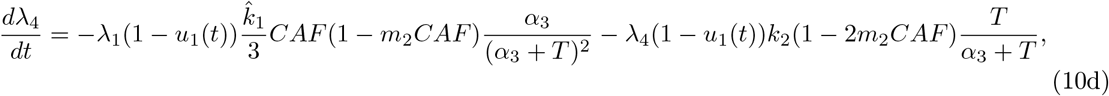

*with λ*_*i*_(*t*_*f*_) = 0, *i* = 1, …, 4. *In addition, u*^∗^ *can be represented by*

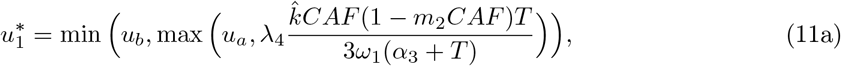

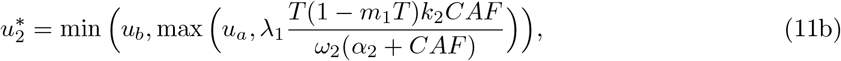

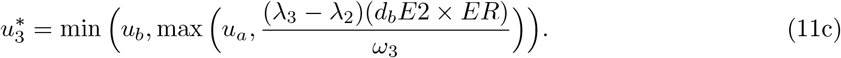

*Proof*. The proof is based on the Pontryagin’s Maximum Principle. We firstly construct the Lagrangian [9] as

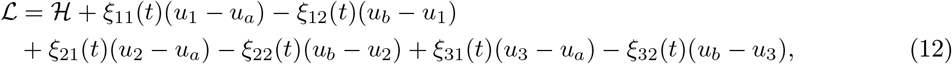

where the Hamiltonian ℋ is defined as

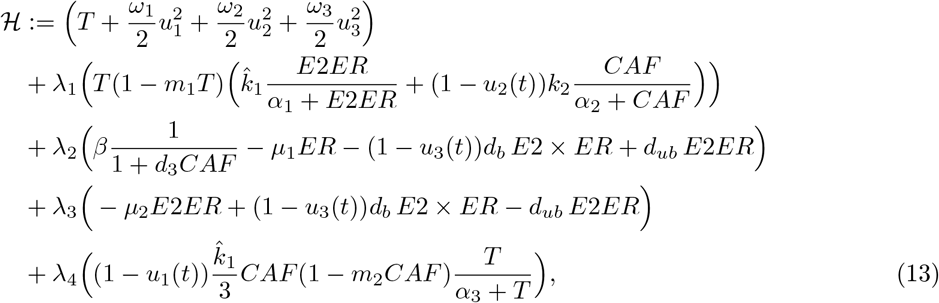

and *ξ*_*i*_(*t*) ≥ 0 are penalty multipliers such that *ξ*_11_(*t*)(*u*_1_−*u*_*a*_) = 0, *ξ*_12_(*t*)(*u*_*b*_−*u*_1_) = 0, *ξ*_21_(*t*)(*u*_2_−*u*_*a*_) = 0, *ξ*_22_(*t*)(*u*_*b*_−*u*_2_) = 0, *ξ*_31_(*t*)(*u*_3_−*u*_*a*_) = 0, *ξ*_32_(*t*)(*u*_*b*_−*u*_3_) = 0 at *u*^∗^. From the Pontryagin’s Maximum Principle, we can derive the adjoint equations as

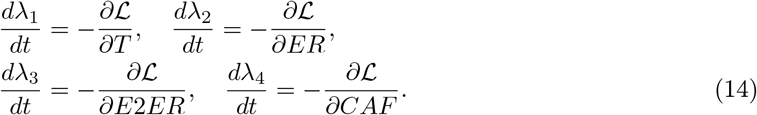

To obtain a formula for the control functions, we differentiate the Hamiltonian with respect to *u*_1_, *u*_2_ and *u*_3_ separately and project them onto the admissible set of controls.

## 3 Results

We firstly consider model (5) and perform model calibration after fixing some model parameters as described in the Material and Methods section. All parameter values are provided in Table 1. We present boxplots of model parameters for three E2 levels in the Appendix B as well. Then, we proceeded with modelling multiple treatment strategies. We compare between constant and optimal treatment aiming to minimize the tumor volume. Treatment is started on the date the first non-zero tumor volume is measured in the experiment for all mice, whereas the final time for simulations is fixed as the last day of the mice experiment.

### 3.1 Constant treatment

We consider five different treatment strategies (Type I, II, III, combination of Type I and III, combination of Type II and III). We perform simulations for model (6) by fixing *ũ*_1_, *ũ*_2_ and *ũ*_3_ as 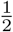 and 0.99 for comparison purposes. We remind the reader that the terms *ũ*_1_ and *ũ*_2_ target the growth of CAFs, and the tumor growth triggered by CAFs and inhibit them, respectively. On the other hand, the term *ũ*_3_ models the inhibition of the binding rate of E2 and ER to represent hormonal treatment.

#### 3.1.1 High dose of E2 (0.5 mg)

We first consider the high E2 condition. We present the simulation results using the parameter values in Table 1. We define the treatment strength as 0.99 and 0.5 in Fig. 4 and Fig. 5 for different ratios of 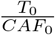, respectively. We observe that tumor grows rapidly in case of no treatment. Treatment of Type I is the least effective one to decrease tumor growth for all mice. The smallest tumor volume is measured when the combination of Type II and Type III are applied together. These observations are independent of the ratio 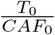 and the strength of treatment. Therefore, inhibiting proliferative effect of CAFs on tumor growth and hormonal treatment was found to be the best treatment option for the high E2 condition. As the population of CAFs increases in size, treatment of Type II becomes less effective; whereas it is the second most successful treatment choice for the highest initial number of CAFs. On the other hand, hormonal treatment is the fourth successful treatment choice for the case *CAF*_0_ = 3*T*_0_, but it becomes the third successful treatment choice when *CAF*_0_ = 0.3*T*_0_. We observe that tumor growth is not correlated with the ratio 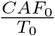. In other words, baseline values of high and low CAF populations lead to bigger tumors than an intermediate number. It seems a nonlinear effect because CAFs influence tumor growth directly and indirectly through ER inhibition. This is non-intuitive and modeling describes such effect.

**Figure 4:**
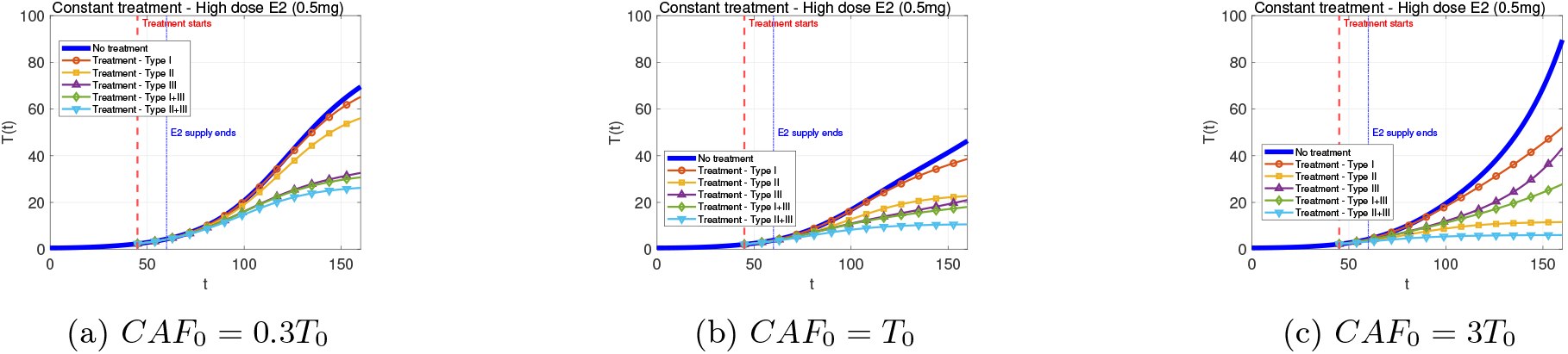
Tumor dynamics *T*(*t*) under constant treatment with a high dose of E2 (0.5 mg) assuming 99% inhibition effect are shown. The blue curve represents the untreated case, while the others correspond to different treatment types: Type I (treatment inhibits the growth of CAFs), Type II (treatment inhibits the tumor growth triggered by CAFs), and Type III (treatment inhibits the binding rate of E2 and ER). Combination treatments (Type I+III and Type II+III) are also shown. The external E2 supply ends at day *t* = 60.

**Figure 5:**
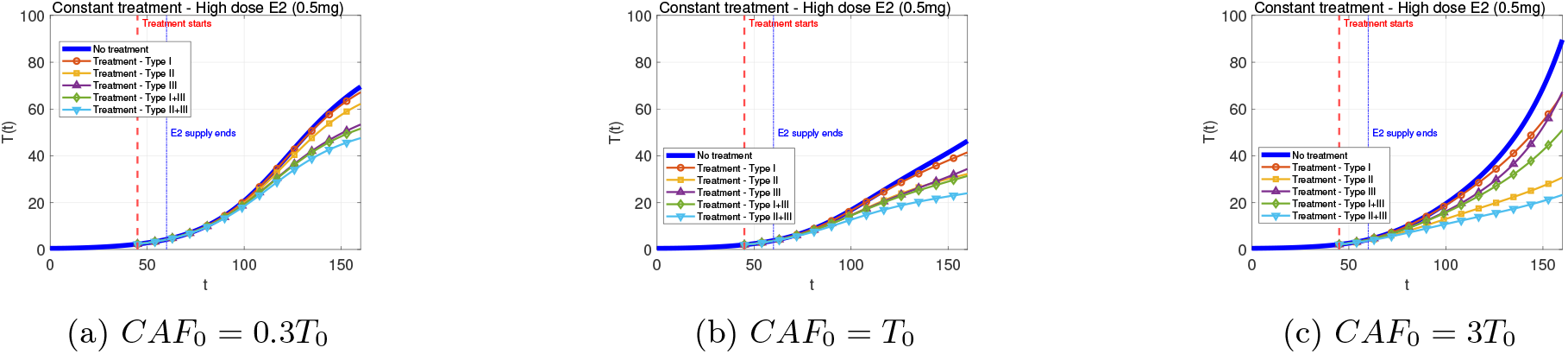
Tumor dynamics *T*(*t*) under constant treatment with a high dose of E2 (0.5 mg) assuming 50% inhibition effect are shown. The blue curve represents the untreated case, while the others correspond to different treatment types: Type I (treatment inhibits the growth of CAFs), Type II (treatment inhibits the tumor growth triggered by CAFs), and Type III (treatment inhibits the binding rate of E2 and ER). Combination treatments (Type I+III and Type II+III) are also shown. The external E2 supply ends at day *t* = 60.

#### 3.1.2 Medium dose of E2 (0.1 mg)

Next, we consider the medium E2 (0.1 mg) condition and display the results for model variables using treatment strength of 0.99 in Fig. 6. Tumor grows up to 30 mm^3^ which is smaller compared to the case of high E2 supply. When we compare different treatment choices, their order with respect to the reduction in final tumor size is equal to the case with high E2 supply. Similarly, the best treatment choice is the combination treatment of Type II and Type III.

**Figure 6:**
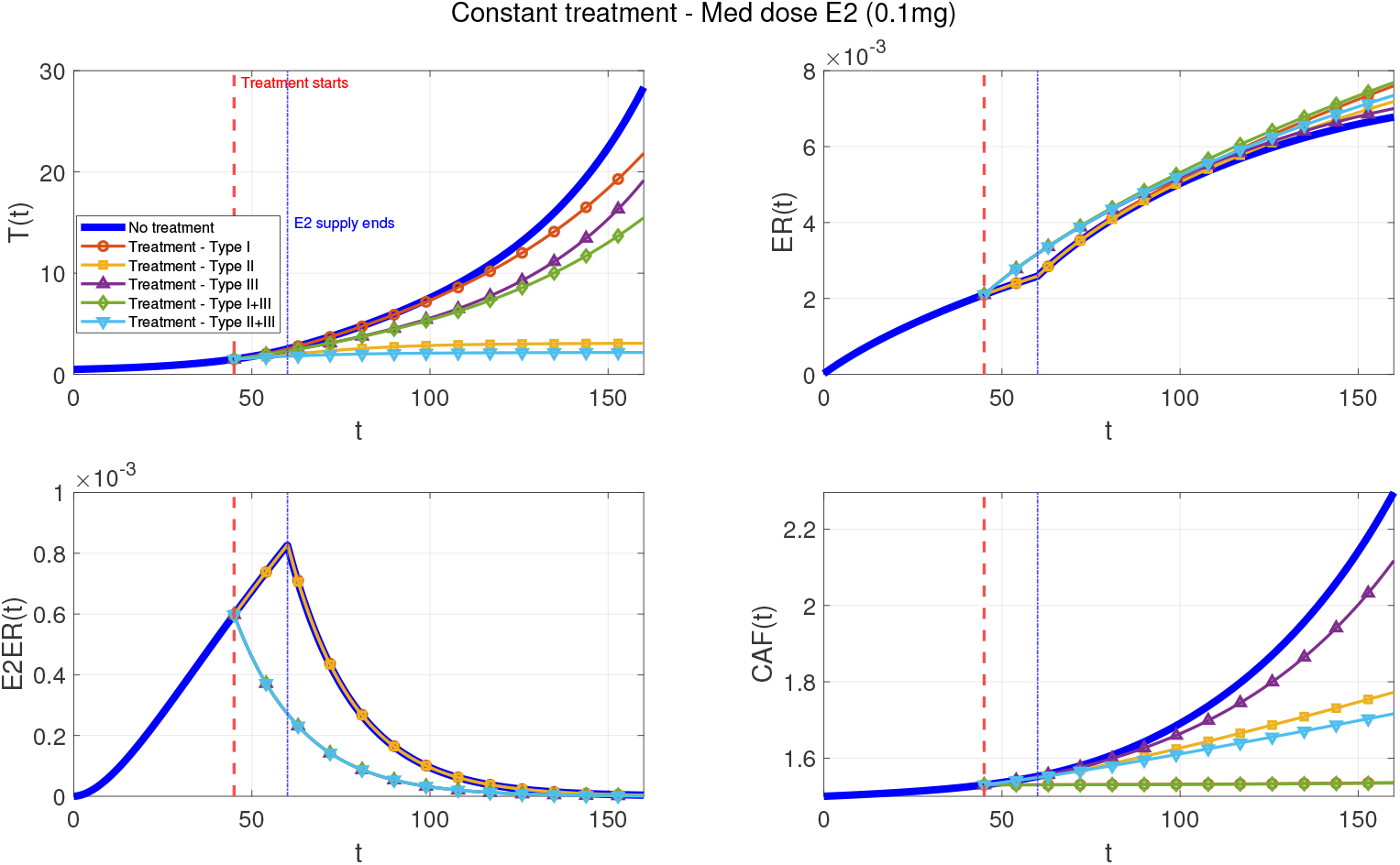
Tumor dynamics *T*(*t*) under constant treatment with a medium dose of E2 (0.1 mg) assuming 99% inhibition effect are shown. The blue curve represents the untreated case, while the others correspond to different treatment types: Type I (treatment inhibits the growth of CAFs), Type II (treatment inhibits the tumor growth triggered by CAFs), and Type III (treatment inhibits the binding rate of E2 and ER). Combination treatments (Type I+III and Type II+III) are also shown. The external E2 supply ends at day *t* = 60. The results correspond to the setting *CAF*_0_ = 3*T*_0_.

#### 3.1.3 Low dose of E2 (0.025 mg)

We proceed with the case of supplying low dose of E2 (0.025 mg) in Fig. 7. tumor growth is even less pronounced due to low dose of E2 supply. The best treatment choice was found as Type II and the combination treatment of Type II and Type III.

**Figure 7:**
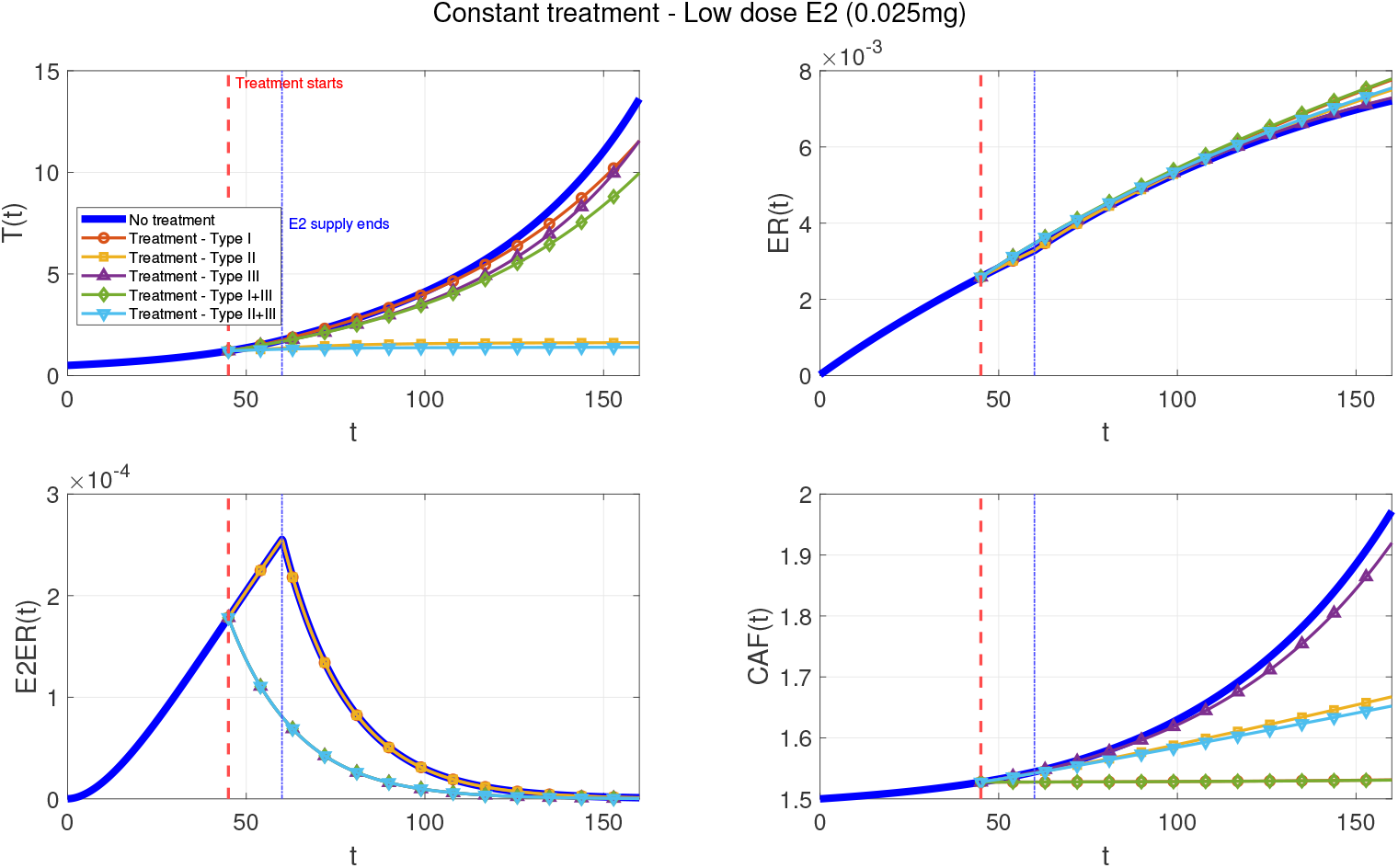
Tumor dynamics *T*(*t*) under constant treatment with a low dose of E2 (0.025 mg) assuming 99% inhibition effect are shown. The blue curve represents the untreated case, while the others correspond to different treatment types: Type I (treatment inhibits the growth of CAFs), Type II (treatment inhibits the tumor growth triggered by CAFs), and Type III (treatment inhibits the binding rate of E2 and ER). Combination treatments (Type I+III and Type II+III) are also shown. The external E2 supply ends at day *t* = 60. The results correspond to the setting *CAF*_0_ = 3*T*_0_.

In the next section, we explore optimal treatment strategies and compare them to the previous constant strategies to find out if unnecessary treatment can be spared.

### 3.2 Optimal treatment

For implementation of the OCP, the box constraints are fixed as *u*_*a*_ = 0 and *u*_*b*_ = 0.99 where *u*_*a*_ and *u*_*b*_ refer to no treatment and the strongest possible treatment, respectively. Here, *u*_*b*_ is not fixed as one since we suppose that no treatment could fully inhibit the corresponding mechanism. As the algorithm to solve the OCP, the forward-backward sweep method with the “greedy” convex combination of the control is used to scan through several combinations of control functions and eliminate stagnation [29, 50]. We refer to the reader to the study [1, Sec. 4.1] for further details on implementation. All data and code are available (see data and code availability part for the details).

#### 3.2.1 High dose of E2 (0.5 mg)

We consider the case when high dose of E2 is supplied in Fig. 8. Similar to the constant treatment, the most effective strategy to decrease tumor volume is combination of Type II and III, while treatment of Type I is the least successful choice. We model the optimal treatment scheduling, so it is found that inhibition terms is not always active (See Fig. 9). Optimal control strategy enables us to eliminate the use of unnecessary drugs to decrease tumor volume.

**Figure 8:**
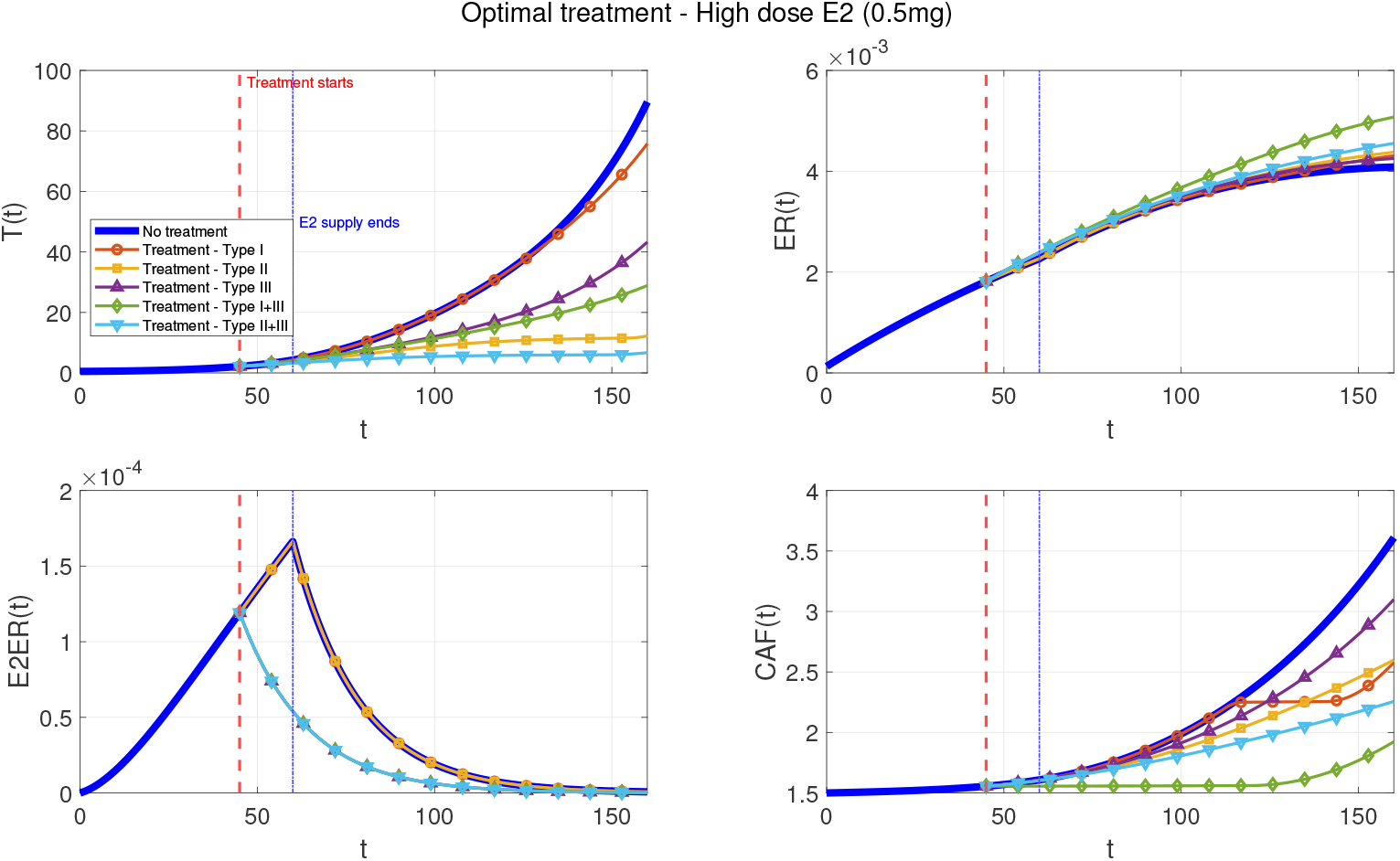
Tumor dynamics *T*(*t*) under optimal treatment with a high dose of E2 (0.5 mg) are shown. The blue curve represents the untreated case, while the others correspond to different treatment types: Type I (treatment inhibits the growth of CAFs), Type II (treatment inhibits the tumor growth triggered by CAFs), and Type III (treatment inhibits the binding rate of E2 and ER). Combination treatments (Type I+III and Type II+III) are also shown. The external E2 supply ends at day *t* = 60. The results correspond to the setting *CAF*_0_ = 3*T*_0_.

**Figure 9:**
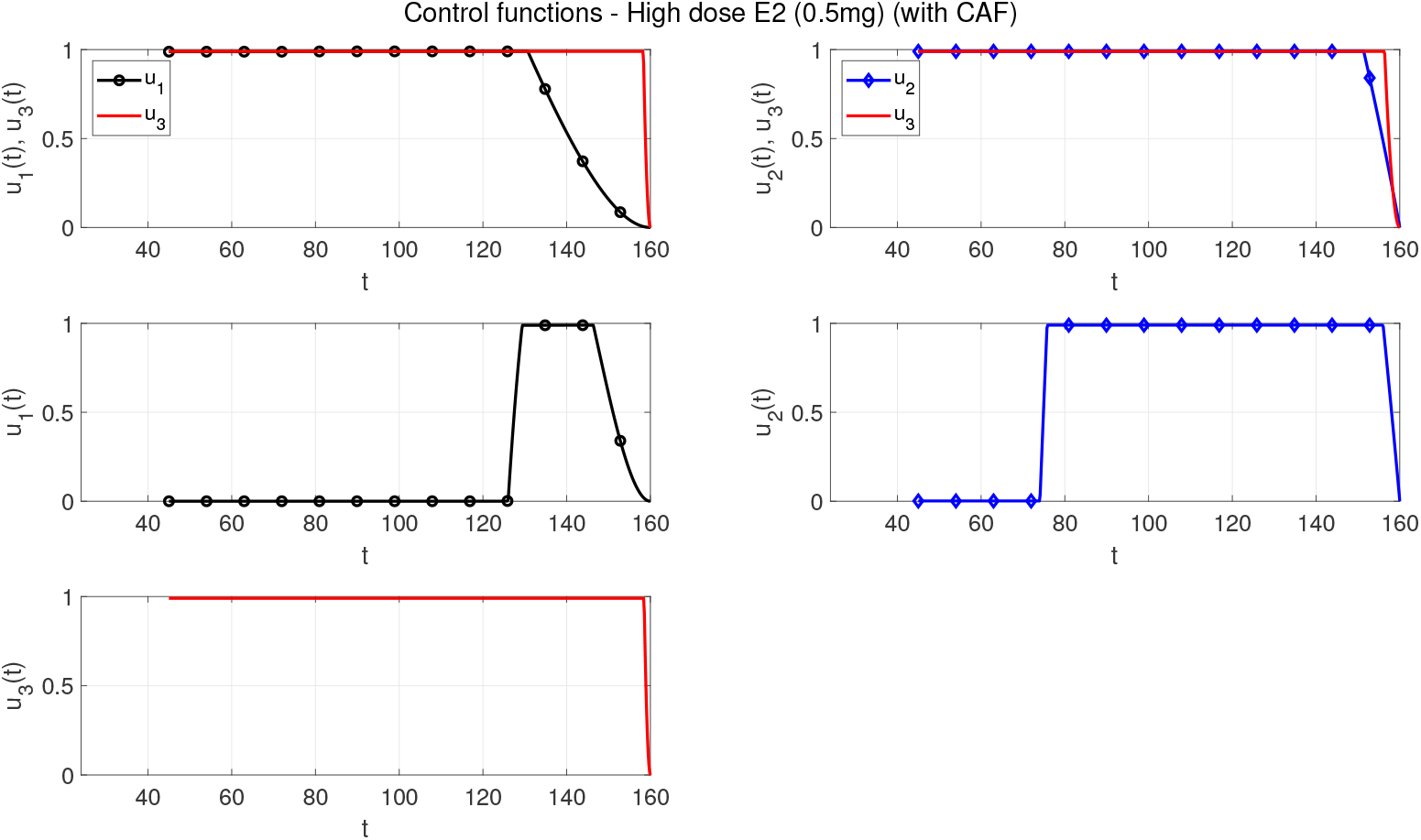
Control functions for high dose of E2 (0.5 mg) are shown for treatments of Type I (treatment inhibits the growth of CAFs), Type II (treatment inhibits the tumor growth triggered by CAFs), Type III (treatment inhibits the binding rate of E2 and ER) and their combinations (Type I+III and Type II+III).

#### 3.2.2 Medium dose of E2 (0.1 mg)

Combination treatment of Type II and Type III was resulted as the highest reduction in tumor volume for medium dose of E2 supply (see Fig. 10), even though the strongest treatment is not applied during simulation (see Fig. 11). Treatment of Type I does not lead to any change in tumor volume.

**Figure 10:**
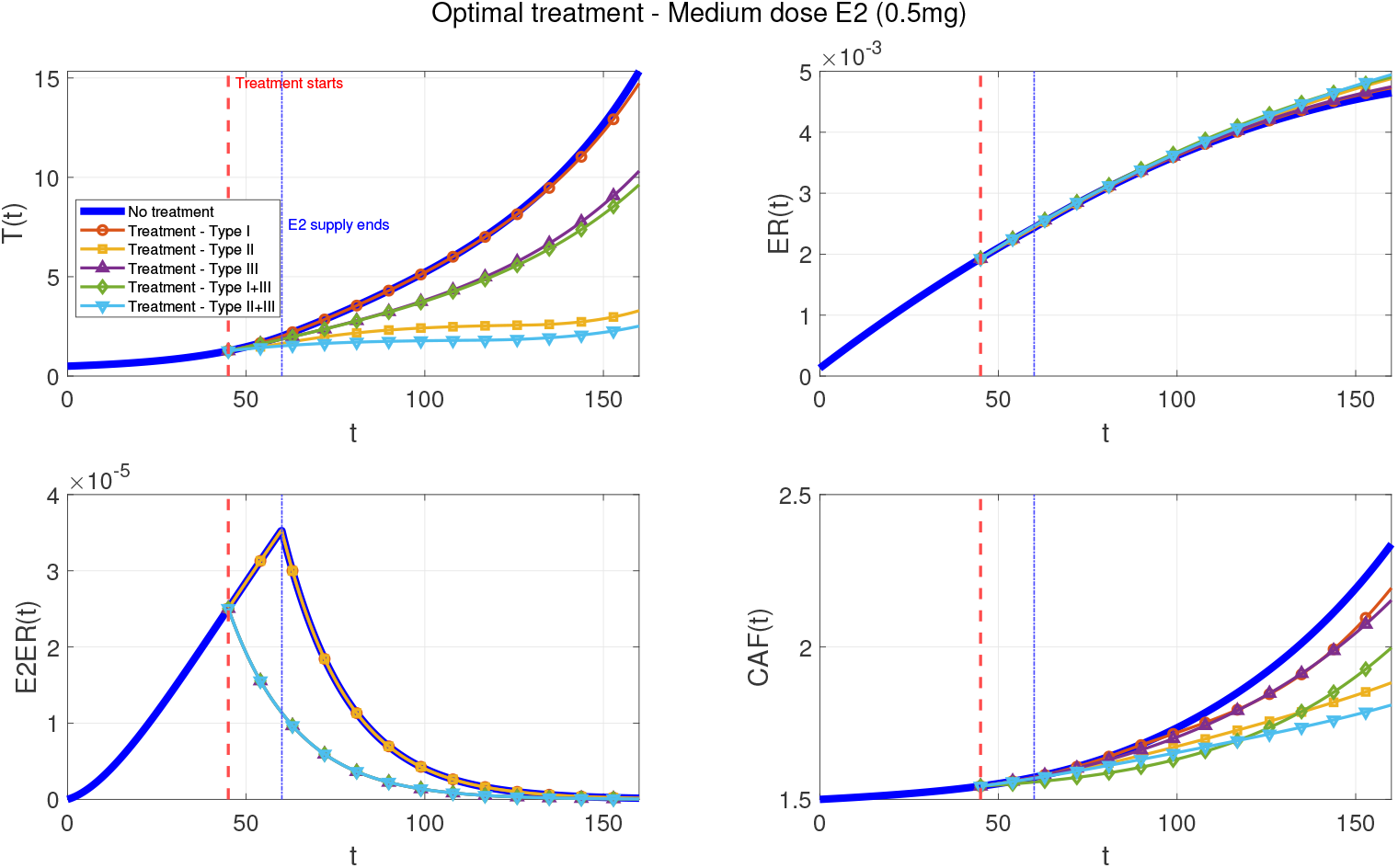
Tumor dynamics *T*(*t*) under optimal treatment with a medium dose of E2 (0.1 mg) are shown. The blue curve represents the untreated case, while the others correspond to different treatment types: Type I (treatment inhibits the growth of CAFs), Type II (treatment inhibits the tumor growth triggered by CAFs), and Type III (treatment inhibits the binding rate of E2 and ER). Combination treatments (Type I+III and Type II+III) are also shown. The external E2 supply ends at day *t* = 60. The results correspond to the setting *CAF*_0_ = 3*T*_0_.

**Figure 11:**
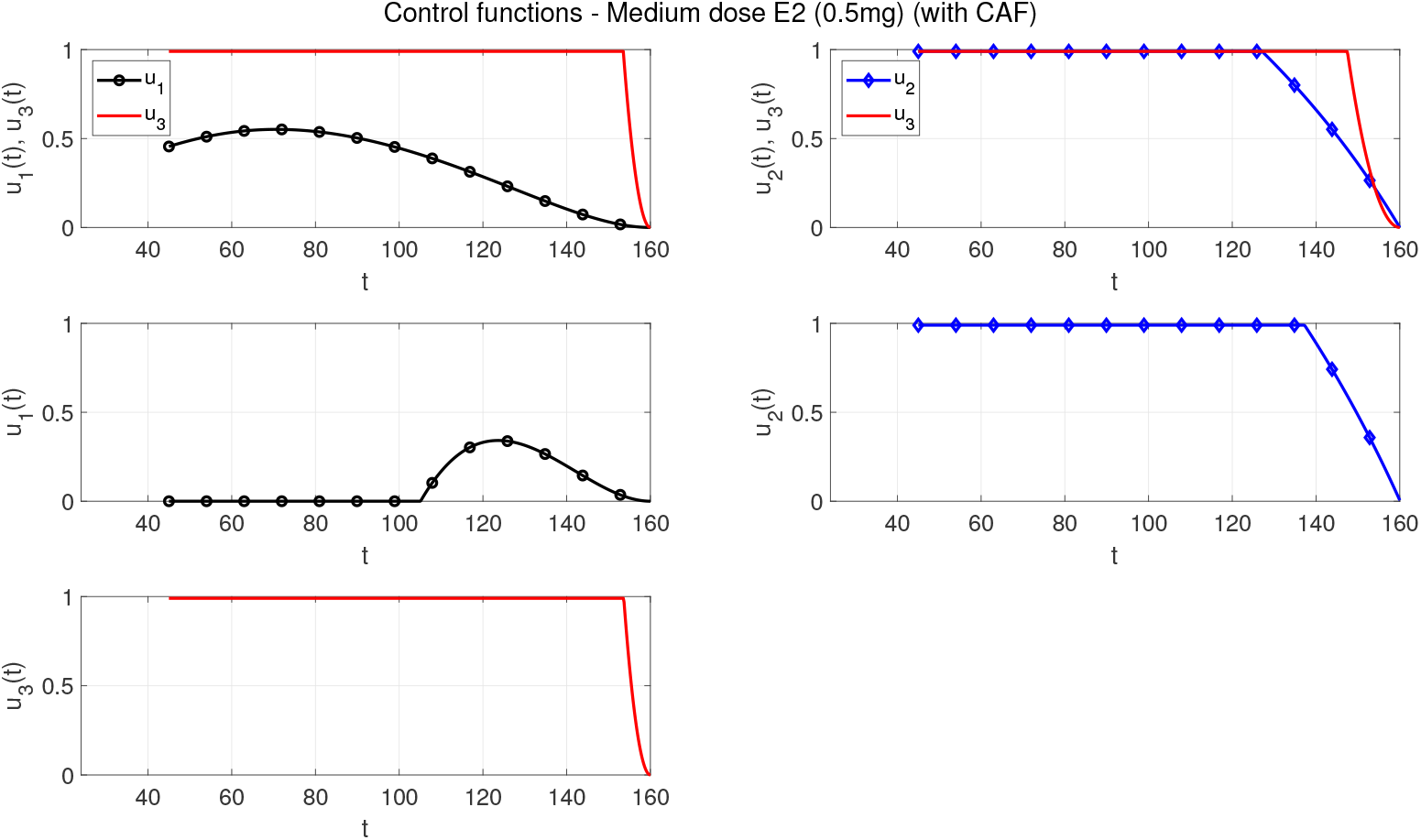
Control functions for medium dose of E2 (0.1 mg) are shown for treatments of Type I (treatment inhibits the growth of CAFs), Type II (treatment inhibits the tumor growth triggered by CAFs), Type III (treatment inhibits the binding rate of E2 and ER) and their combinations (Type I+III and Type II+III).

#### 3.2.3 Low dose of E2 (0.025 mg)

In case of low dose of E2, combination treatment of Type II and Type III was found as the most effective option in reducing tumor growth (see Fig. 12). If we compare the optimal treatment schedules, different treatments should act longer than their corresponding plots for high dose of E2 (see Fig. 13).

**Figure 12:**
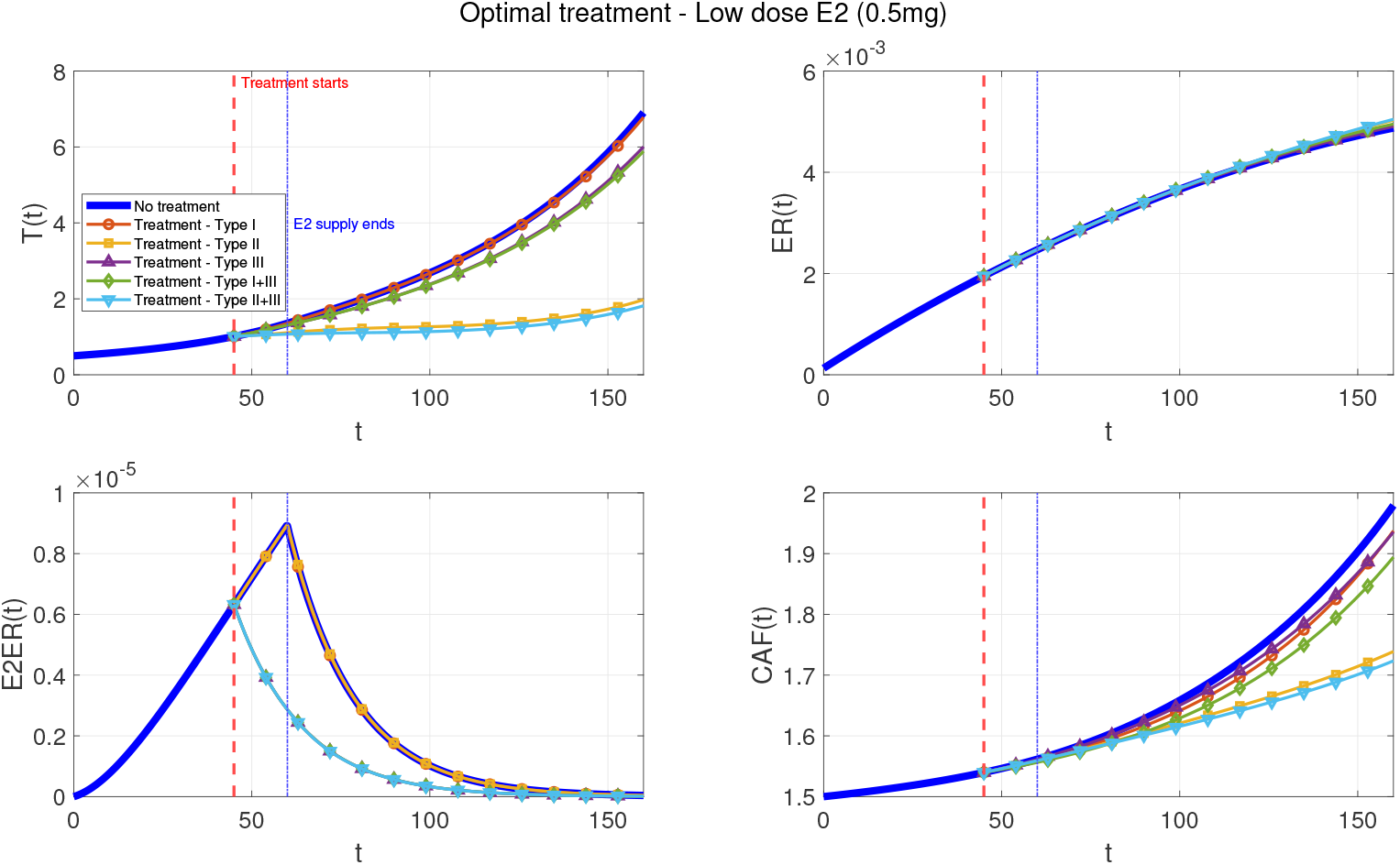
Tumor dynamics *T*(*t*) under optimal treatment with a low dose of E2 (0.025 mg) are shown. The blue curve represents the untreated case, while the others correspond to different treatment types: Type I (treatment inhibits the growth of CAFs), Type II (treatment inhibits the tumor growth triggered by CAFs), and Type III (treatment inhibits the binding rate of E2 and ER). Combination treatments (Type I+III and Type II+III) are also shown. The external E2 supply ends at day *t* = 60. The results correspond to the setting *CAF*_0_ = 3*T*_0_.

**Figure 13:**
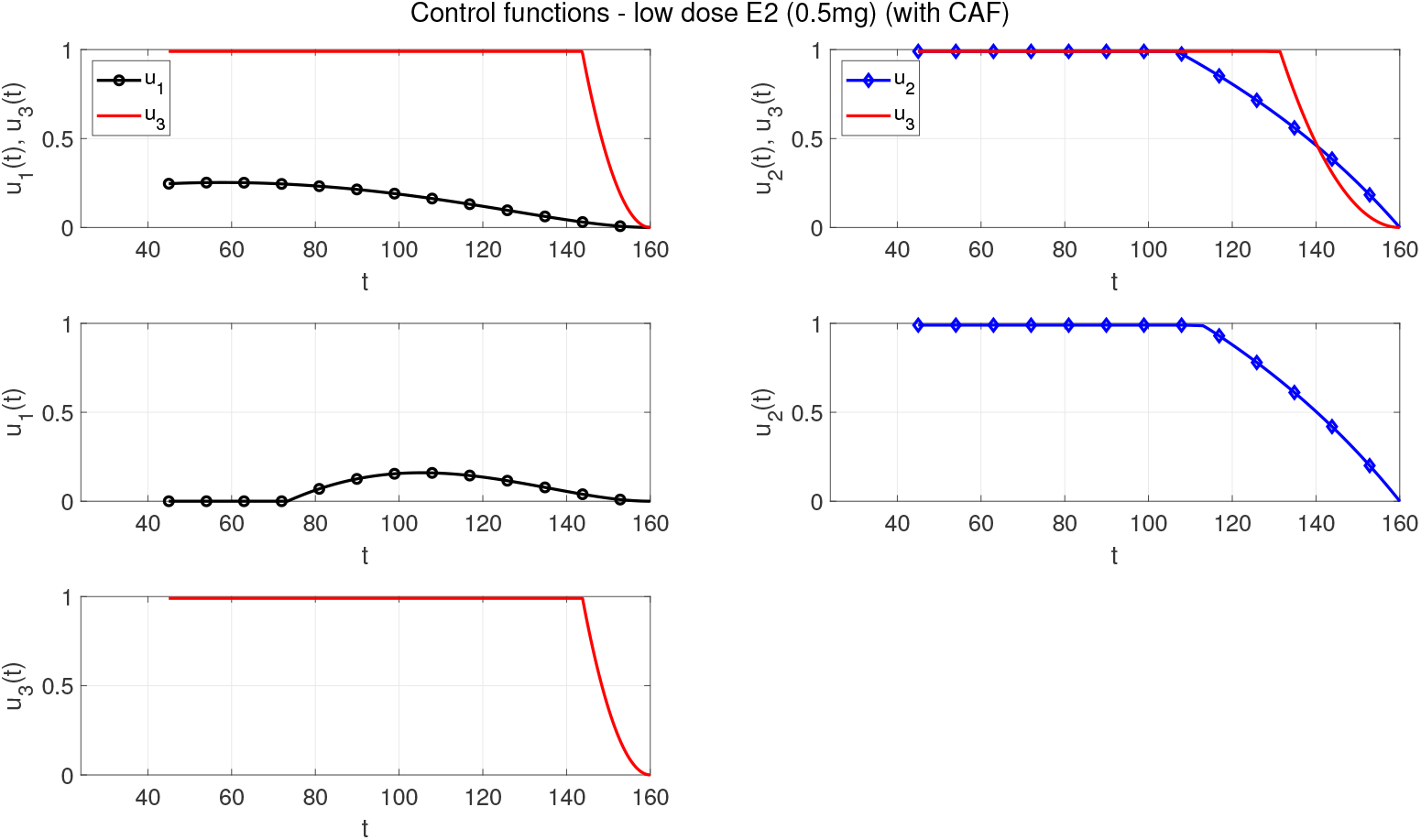
Control functions for low dose of E2 (0.025 mg) are shown for treatments of Type I (treatment inhibits the growth of CAFs), Type II (treatment inhibits the tumor growth triggered by CAFs), Type III (treatment inhibits the binding rate of E2 and ER) and their combinations (Type I+III and Type II+III).

We compare percentage reduction of the final tumor volume across different treatment choices for three different E2 conditions in case of different ratios of 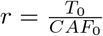 in Table 2. For all E2 conditions, the best treatment choice is combination treatment of Type II and Type III. Constant treatment with 99% inhibition results in highest tumor reduction than 50% inhibition, as expected. For high E2 condition, treatment of Type III leads to the same reduction for all ratios of *r*. There is a positive correlation between the percentage reduction and the ratio *r* for all treatment types, except Type III. It means that as the initial population of CAFs is less pronounced, then percentage reduction decreases. For medium and low E2 conditions, there is a negative correlation between the ratio *r* and percentage reduction for treatment Type III and combination treatment of Type I and III. When presence of CAFs in the beginning of the experiment is less strong, contribution of hormonal treatment is revealed for both constant and optimal treatment. Optimal treatment leads to the smallest reduction than the constant treatment Type I for all E2 conditions. On the other hand, optimal treatment results in the smallest percentage reduction for low E2 condition and treatment Type II, as well. For all other cases, optimal treatment scheduling is a very strong alternative against constant treatment.

**Table 2:** Percentage reduction of the final tumor volume across different treatments (Type I: treatment inhibits the growth of CAFs, Type II: treatment inhibits the tumor growth triggered by CAFs, Type III: treatment inhibits the binding rate of E2 and ER).

| Treatment type | 99% inhibition |  |  |  | 50% inhibition |  |  |  | Optimal treatment |  |  |  |
| --- | --- | --- | --- | --- | --- | --- | --- | --- | --- | --- | --- | --- |
| $r = \frac{T_0}{CAF_0}$ | $\frac{1}{3}$ | 1 | 2 | 4 | $\frac{1}{3}$ | 1 | 2 | 4 | $\frac{1}{3}$ | 1 | 2 | 4 |
| *High dose of E2* |  |  |  |  |  |  |  |  |  |  |  |  |
| Type I | 41.83 | 16.73 | 10.17 | 5.59 | 25.97 | 10.50 | 5.94 | 3.15 | 15.38 | 0.49 | 0 | 0 |
| Type II | 87.07 | 51.05 | 30.35 | 16.76 | 65.69 | 30.41 | 17.37 | 9.21 | 86.30 | 44.63 | 24.14 | 12.55 |
| Type III | 51.69 | 54.66 | 54.37 | 52.58 | 24.88 | 25.78 | 25.28 | 22.82 | 51.69 | 54.66 | 54.37 | 52.58 |
| Type I-III | 68.96 | 61.03 | 58.24 | 54.95 | 43.09 | 32.26 | 29.04 | 25.00 | 67.76 | 57.37 | 54.95 | 52.69 |
| Type II-III | 93.17 | 77.14 | 68.06 | 60.55 | 73.98 | 48.18 | 37.46 | 29.53 | 92.51 | 75.65 | 66.80 | 59.62 |
| *Medium dose of E2* |  |  |  |  |  |  |  |  |  |  |  |  |
| Type I | 21.13 | 7.38 | 4.94 | 3.40 | 11.97 | 4.03 | 2.15 | 1.32 | 3.97 | 0 | 0 | 0 |
| Type II | 82.98 | 45.92 | 26.98 | 15.69 | 60.25 | 26.89 | 15.06 | 7.44 | 78.53 | 37.05 | 18.45 | 7.51 |
| Type III | 32.73 | 45.20 | 50.50 | 53.48 | 16.36 | 22.27 | 24.01 | 25.22 | 32.73 | 45.20 | 50.49 | 53.48 |
| Type I-III | 44.67 | 48.26 | 52.30 | 54.66 | 25.67 | 25.14 | 25.97 | 26.41 | 37.24 | 45.38 | 50.55 | 53.49 |
| Type II-III | 88.04 | 69.65 | 63.57 | 60.41 | 66.35 | 42.75 | 35.53 | 30.88 | 83.60 | 60.84 | 56.89 | 56.26 |
| *Low dose of E2* |  |  |  |  |  |  |  |  |  |  |  |  |
| Type I | 13.30 | 3.37 | 2.31 | 1.09 | 7.18 | 1.87 | 1.03 | 0.6 | 1.78 | 0 | 0 | 0 |
| Type II | 81.24 | 43.17 | 25.21 | 13.76 | 57.77 | 24.93 | 13.80 | 7.22 | 71.36 | 18.06 | 6.67 | 2.28 |
| Type III | 13.05 | 25.73 | 35.52 | 43.74 | 6.58 | 12.99 | 17.64 | 21.34 | 13.05 | 25.73 | 35.52 | 43.74 |
| Type I-III | 23.79 | 27.84 | 36.55 | 44.32 | 13.09 | 14.36 | 18.41 | 21.84 | 14.77 | 25.75 | 35.53 | 43.75 |
| Type II-III | 83.52 | 57.63 | 51.51 | 51.39 | 60.48 | 34.69 | 28.96 | 27.18 | 73.68 | 36.42 | 38.35 | 44.64 |

We split the cost functional 𝒥 defined in Eq. (7) into two parts and compare the contribution of tumor size (𝒥_1_) and treatment (𝒥_2_) to its value as follows:

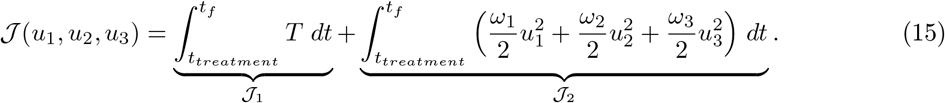

We present the values of 𝒥_1_, 𝒥_2_ and 𝒥 under high, medium, and low E2 conditions when constant and optimal treatment are applied in case of 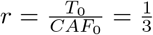 in Table 3. Our model reveals that tumor size decreases as we increase the strength of treatment for all ratios of 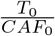. For high E2 condition, the smallest value of 𝒥 is obtained for constant treatment of Type I with strength *s* = 0.99 due to the smallest total tumor volume (𝒥_1_) measured during the simulation. For other treatment choices, the smallest value of 𝒥 is derived for optimal treatment due to effect of optimal treatment. For medium E2 condition, constant treatment with *s* = 0.5 leads to the smallest value of 𝒥 due to weaker treatment. For other treatment choices, the smallest value of 𝒥 is achieved via optimal treatment even though a smaller tumor volume is measured for constant treatment. As different from the other cases, for low E2 condition, optimal treatment results in the smallest value of 𝒥 due to optimal treatment scheduling (J_2_).

**Table 3:** Comparison of the values of 𝒥_1_, 𝒥_2_ and 𝒥 under high, medium, and low E2 conditions when constant and optimal treatment are applied in case of 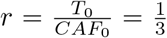 (Type I: treatment inhibits the growth of CAFs, Type II: treatment inhibits the tumor growth triggered by CAFs, Type III: treatment inhibits the binding rate of E2 and ER).

| Treatment type | 99% inhibition |  |  | 50% inhibition |  |  | Optimal treatment |  |  |
| --- | --- | --- | --- | --- | --- | --- | --- | --- | --- |
| *High dose of E2* | $\mathcal{J}_1$ | $\mathcal{J}_2$ | $\mathcal{J}$ | $\mathcal{J}_1$ | $\mathcal{J}_2$ | $\mathcal{J}$ | $\mathcal{J}_1$ | $\mathcal{J}_2$ | $\mathcal{J}$ |
| Type I | 2503.133 | 56.356 | 2559.489 | 2812.038 | 14.375 | 2826.413 | 3075.849 | 10.518 | 3086.368 |
| Type II | 930.614 | 56.356 | 986.97 | 1645.881 | 14.375 | 1660.256 | 931.889 | 40.281 | 972.170 |
| Type III | 1785.100 | 56.356 | 1841.456 | 2559.780 | 14.375 | 2574.155 | 1785.117 | 55.764 | 1840.881 |
| Type I-III | 1465.592 | 112.712 | 1578.304 | 2256.860 | 28.750 | 2285.610 | 1475.366 | 101.026 | 1576.392 |
| Type II-III | 570.496 | 112.712 | 683.208 | 1327.697 | 28.750 | 1356.447 | 573.108 | 108.553 | 681.661 |
| *Medium dose of E2* |  |  |  |  |  |  |  |  |  |
| Type I | 654.606 | 56.356 | 710.962 | 685.806 | 14.375 | 700.181 | 712.679 | 1.395 | 714.074 |
| Type II | 259.406 | 56.356 | 315.761 | 415.477 | 14.375 | 429.852 | 265.993 | 49.279 | 315.271 |
| Type III | 521.813 | 56.356 | 578.169 | 624.419 | 14.375 | 638.794 | 521.815 | 53.860 | 575.676 |
| Type I-III | 480.652 | 112.712 | 593.364 | 594.102 | 28.750 | 622.852 | 504.634 | 63.004 | 567.637 |
| Type II-III | 197.364 | 112.712 | 310.076 | 363.234 | 28.750 | 391.984 | 206.543 | 97.770 | 304.312 |
| *Low dose of E2* |  |  |  |  |  |  |  |  |  |
| Type I | 341.614 | 56.356 | 397.970 | 351.567 | 14.375 | 365.942 | 359.469 | 0.501 | 359.970 |
| Type II | 141.203 | 56.356 | 197.559 | 218.141 | 14.375 | 232.516 | 153.839 | 42.320 | 196.159 |
| Type III | 322.332 | 56.356 | 378.687 | 342.338 | 14.375 | 356.713 | 322.332 | 49.992 | 372.324 |
| Type I-III | 305.162 | 112.712 | 417.874 | 332.284 | 28.750 | 361.034 | 319.260 | 51.807 | 371.067 |
| Type II-III | 127.675 | 112.712 | 240.386 | 206.680 | 28.750 | 235.430 | 141.676 | 85.724 | 227.399 |

## 4 Discussion

Here, we show a mathematical model representing the dynamics of ER signalling and the activity of CAFs to reveal their combined effect on the progression and treatment response of ER+ breast cancer. Utilizing the experimental data generated by Reid et al. [42], we derived calibrated nonlinear ordinary differential equations and developed optimal control theory to establish a framework that provides new insights into the role of CAFs to facilitate tumor growth dynamics, endocrine sensitivity and optimal therapeutic strategy. Our model reinforces the experimental data exhibiting the role of CAFs in regulating the estrogen-independent growth dynamics of luminal breast cancer [5, 42, 4].

Consistent with in vivo findings where co-transplantation of CAFs restored tumor growth following the removal of estrogen [42], low-estrogen conditions elevated tumor persistence with the presence of CAFs, as revealed by the simulations. The effects observed in simulations are due to the paracrine effect of CAFs modulating ER*α* transcription, which is evident with prior findings whereby IL-6 secreted by CAFs can modulated ER*α* signalling and treatment resistance in breast cancer [5]. The parameter calibration of estrogen dosing (high, medium, and low) performed in our study further demonstrated that the dependency to the presence of CAFs increases as estrogen dosing decreases. This aligns with the notion that targeting CAFs in the tumor stroma can be a critical strategy to overcome hormone-deprivation resistance, complementing existing endocrine therapies [17, 41, 5].

The simulations revealed that inhibition of the proliferative effect of CAFs to tumor growth (Type II) and combination of Type II and Type III which also interferes with E2ER binding exhibited the largest reductions in tumor volume with all estrogen dosing. This finding is supported with previous studies wherein the targeting of signalling pathways, namely SDF-1/CXCR4 and TGF-B1 [23], related to the maintenance of CAF phenotype, can decrease tumor progression and therapeutic efficiency. On the other hand, Type I, a direct inhibition of the proliferation of CAFs alone, or Type III, an interference with ER binding, resulted in limited efficiency, especially under low estrogen dosing, suggesting that monotherapies when compared to combination therapy can offer less effective therapeutic outcome in disrupting stromal-epithelial interactions. Our approach, hence, demonstrates the rationale behind using CAF-related combination strategy in ER+ breast cancer, which previous mathematical model-based studies have generally focused heterotypic interactions between tumor-immune cells [10, 21, 11], while few mathematical models explicitly showed CAF-driven endocrine resistance. We observed from simulation results that there is a positive correlation between the percentage reduction of the final tumor volume and the ratio 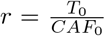 for all treatment types, except Type III. It means that as the initial population of CAFs is less pronounced, then percentage reduction of the final tumor volume is more visible. For medium and low E2 conditions, there is a negative correlation between the ratio *r* and percentage reduction of the final tumor volume for treatment Type III and combination treatment of Type I and III. When presence of CAFs in the beginning of the experiment is less strong, contribution of hormonal treatment is revealed for both constant and optimal treatment more.

We also showed time dependent effect of the interference of CAF-targeting and endocrine therapy in effecting tumor burden from the perspective of an optimal control theory. We derived optimal control schedules that exhibit the duration for the inactivity of treatment, suggesting that avoiding overtreatment wherein continues maximal inhibition is not necessary. This is supported with earlier studies where adaptive or intermittent dosing schedules in oncology could exhibit balanced efficacy and toxicity [44, 21, 9]. Critically, optimal control strategies offered by our model can potentially provide clinical guidance for therapy design, which may be implemented in digital twin approaches as proposed by Wu et al. [54].

While the critical quantitative features of CAF-tumor interactions have been revealed in this study, a number of limitations should be addressed in future work. First, functional and phenotypic heterogeneity of CAFs [4] should be considered in the future as the current work treated CAFs as a homogenous population. Multiple CAF subpopulations can be included into a model framework to refine treatment modalities for overcoming the endocrine resistance. Second, experimental validations for the pharmacodynamics features of CAF-targeted agents including the drug-specific parameters will be important for translational application. Third, further validation, beyond current in vivo xenograft data, using clinical dataset would improve clinical relevance and patient-relevant power of the current model.

Overall, our findings highlight the key role of CAFs in mediating endocrine resistance and tumor growth dynamics in ER+ breast cancer. By taking together mechanistic modelling, experimental data, and optimal control theory, we present a mathematical model for CAF-targeted treatment strategies that could help to improve existing approaches for endocrine therapies.

## A Appendix 1

### Structural identifiability of Model (1)

~~~
***************************************
* RESULTS OF IDENTIFIABILITY ANALYSIS *
***************************************
=> THE MODEL IS STRUCTURALLY GLOBALLY IDENTIFIABLE
Structurally globally identifiable parameters:
   x20
    db
 betaa
   x30
alpha1
k1_hat
Structurally locally identifiable parameters:
 []
Structurally non-identifiable parameters:
 []
~~~

### Structural identifiability of Model (5)

~~~
***************************************
* RESULTS OF IDENTIFIABILITY ANALYSIS *
***************************************
=> THE MODEL IS STRUCTURALLY GLOBALLY IDENTIFIABLE
Structurally globally identifiable parameters:
    d3
alpha3
    k2
alpha2
Structurally locally identifiable parameters:
 []
Structurally non-identifiable parameters:
 []
~~~

## B Appendix 2

### Boxplots for the model parameters

**Figure 14:**
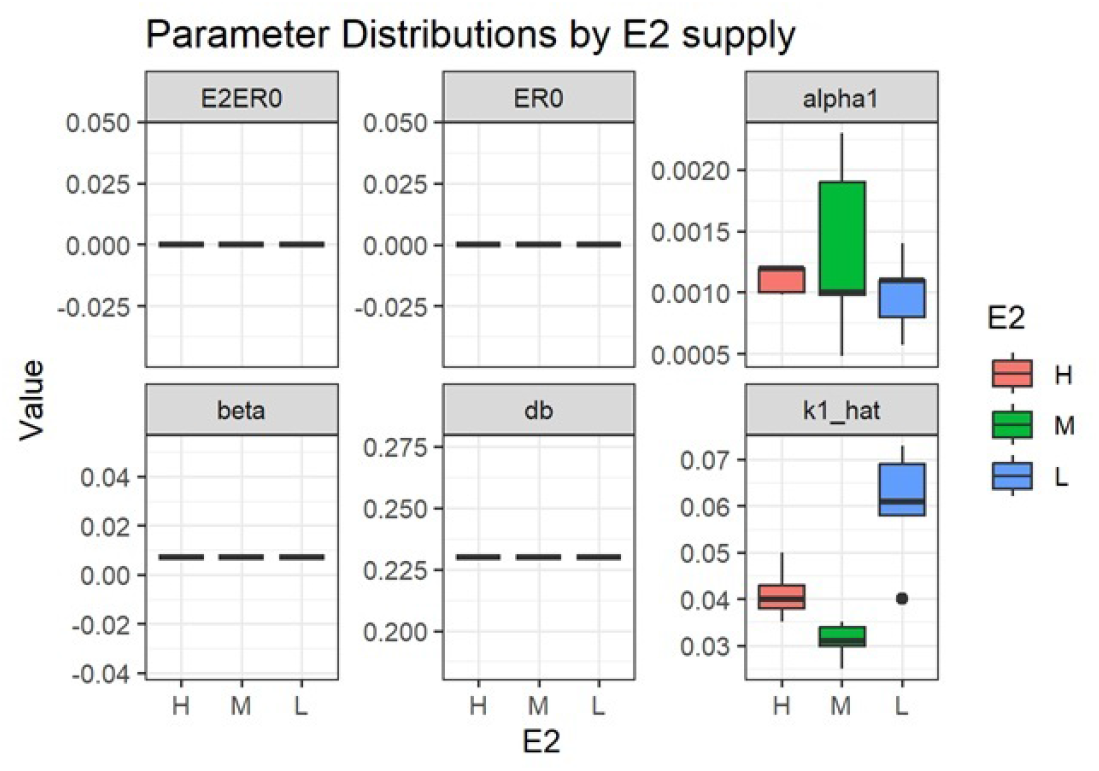
Box plot of the parameters in Model (1).

**Figure 15:**
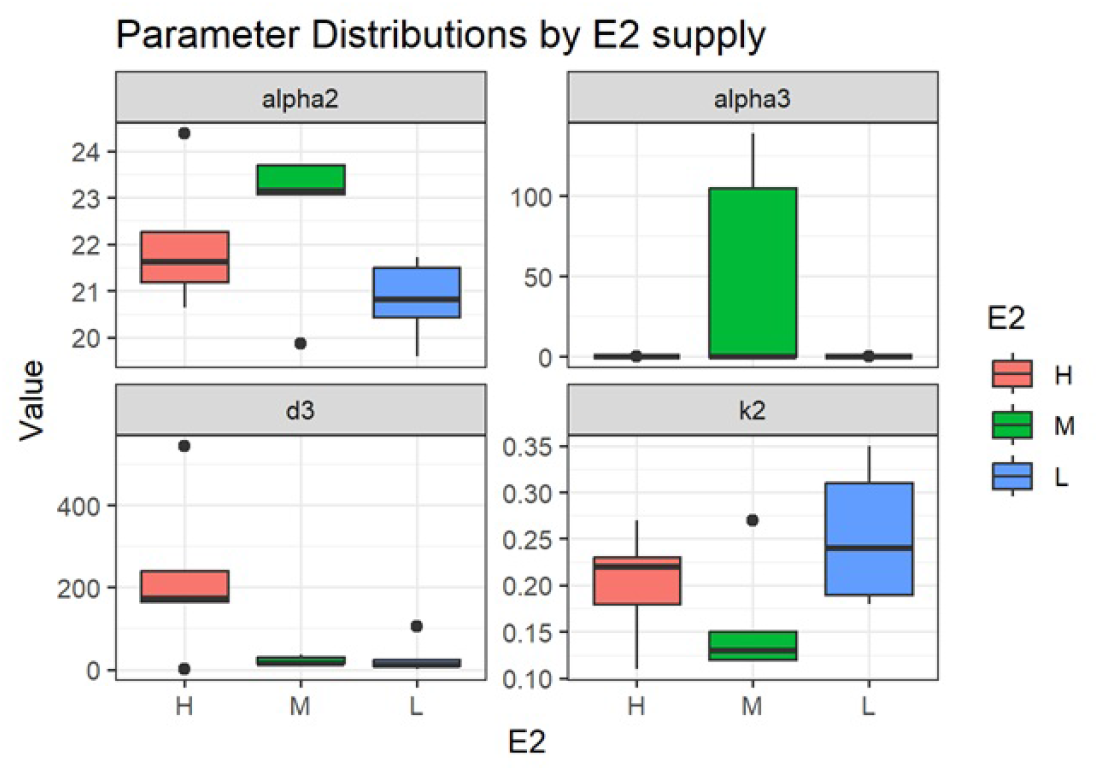
Box plot of the parameters in Model (5).

### Plots for individual fits

**Figure 16:**
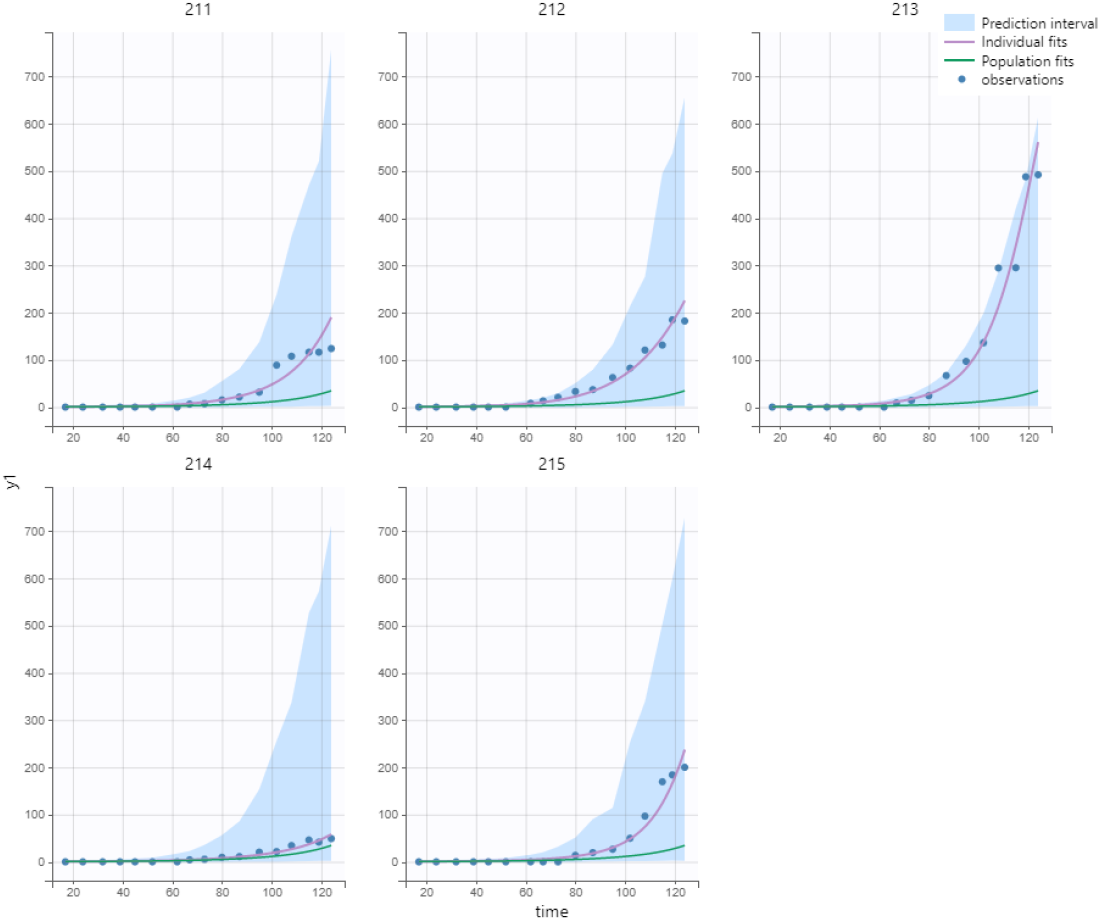
Simulation results for *T*(*t*) in case of high E2 level in Model (1).

**Figure 17:**
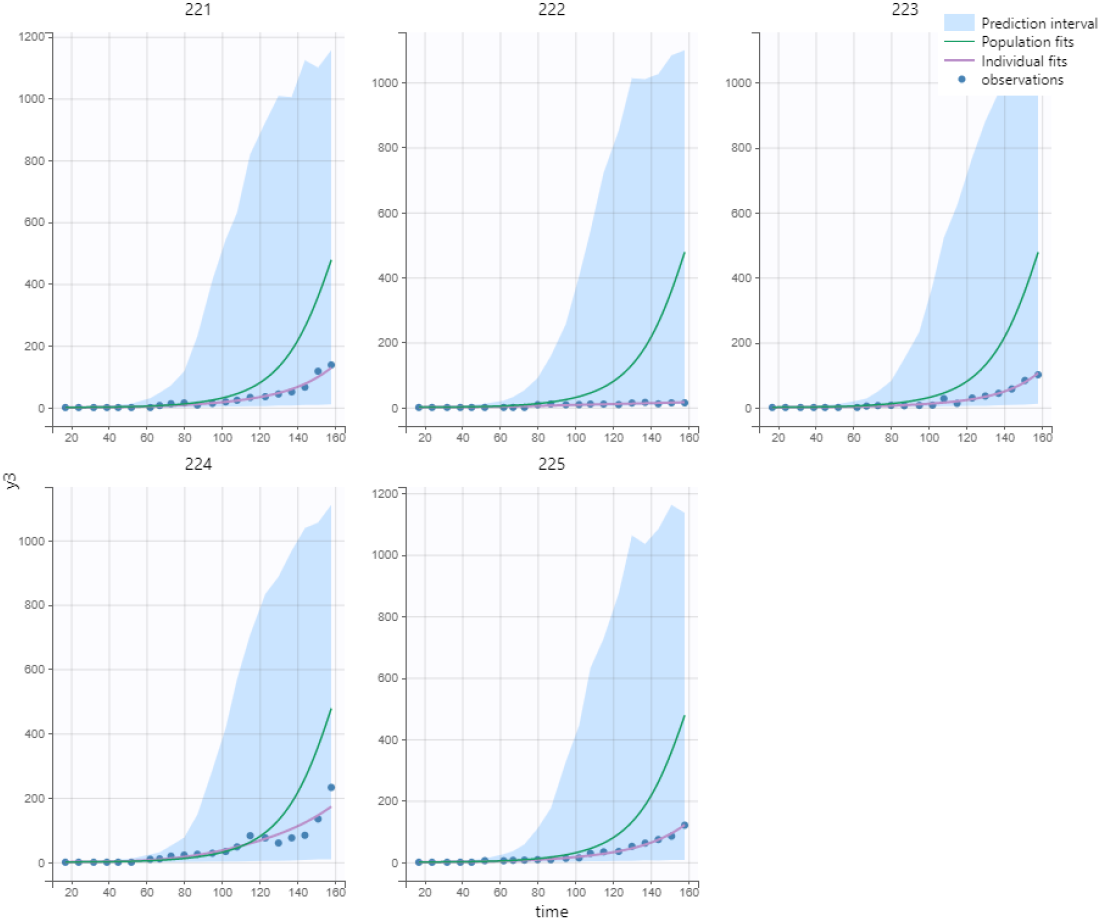
Simulation results for *T*(*t*) in case of medium E2 level in Model (1).

**Figure 18:**
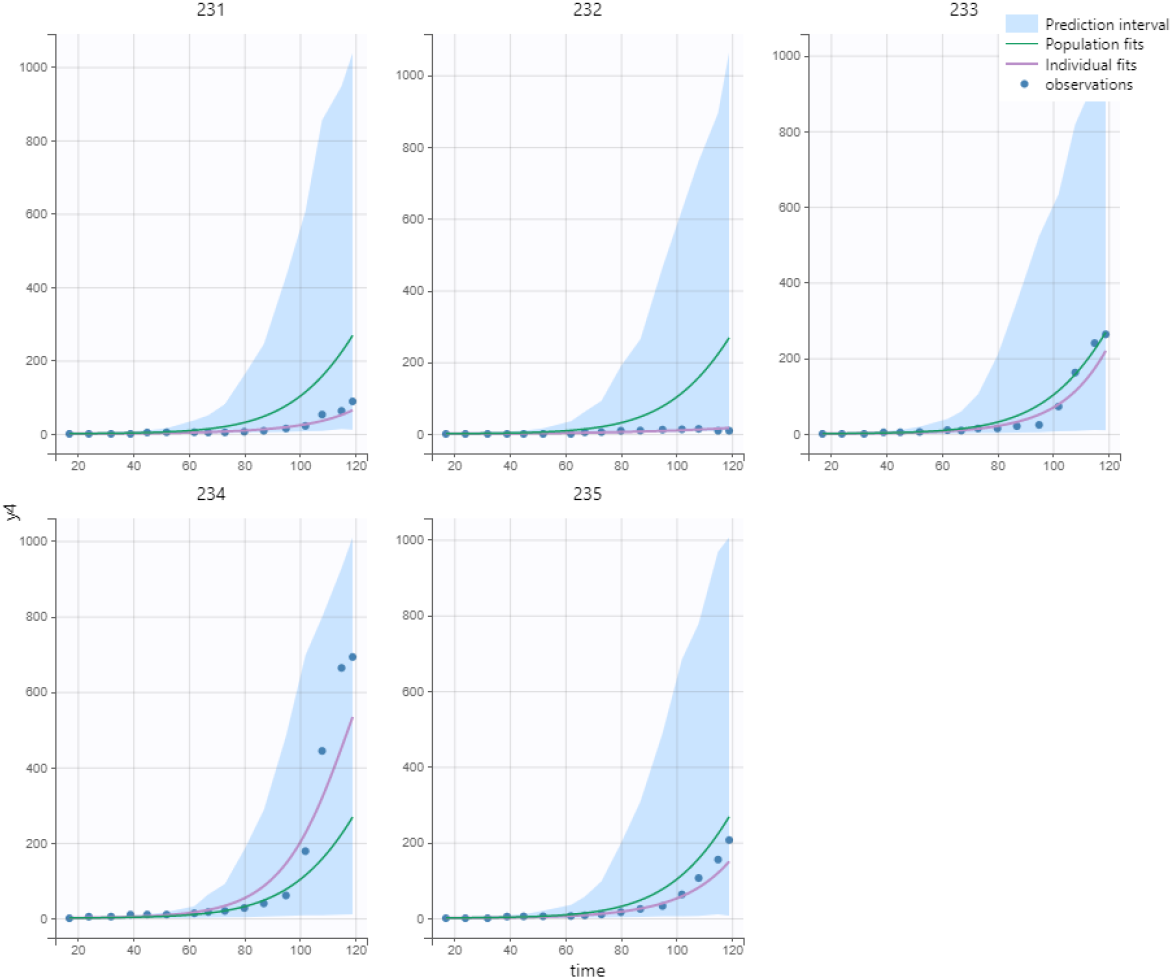
Simulation results for *T*(*t*) in case of low E2 level in Model (1).

**Figure 19:**
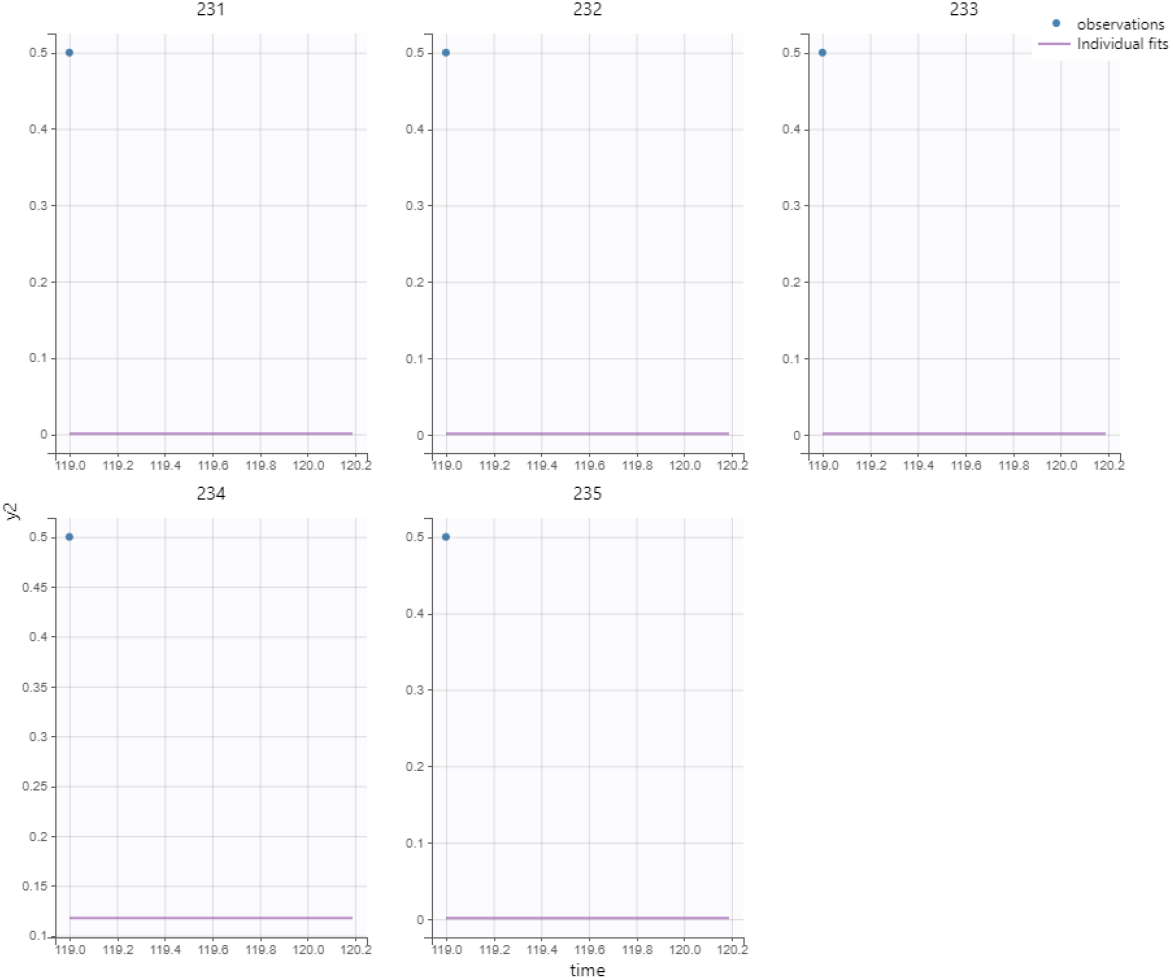
Model calibration for *ER*(*t*) in case of high E2 level in Model (1).

**Figure 20:**
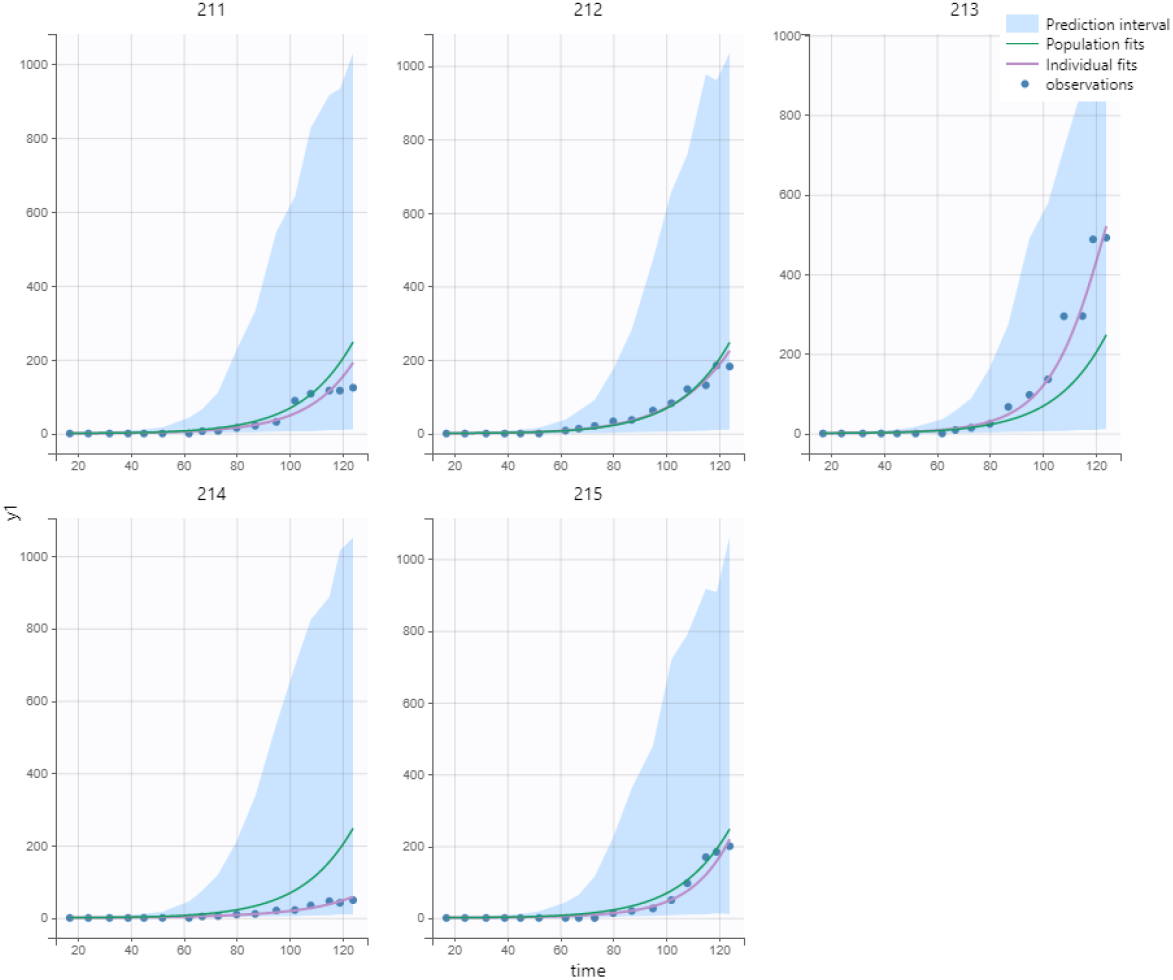
Simulation results for *T*(*t*) in case of high E2 level in Model (5).

**Figure 21:**
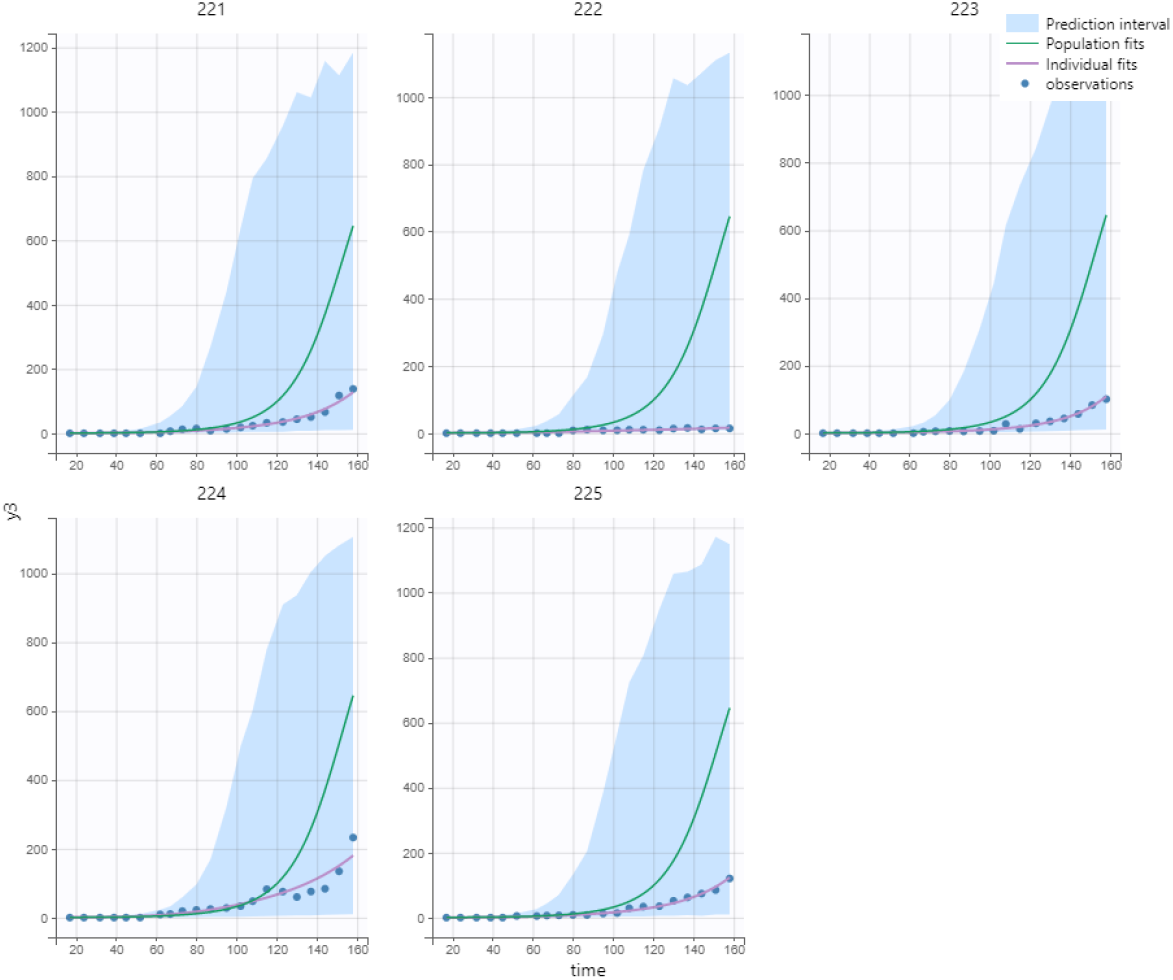
Simulation results for *T*(*t*) in case of medium E2 level in Model (5).

**Figure 22:**
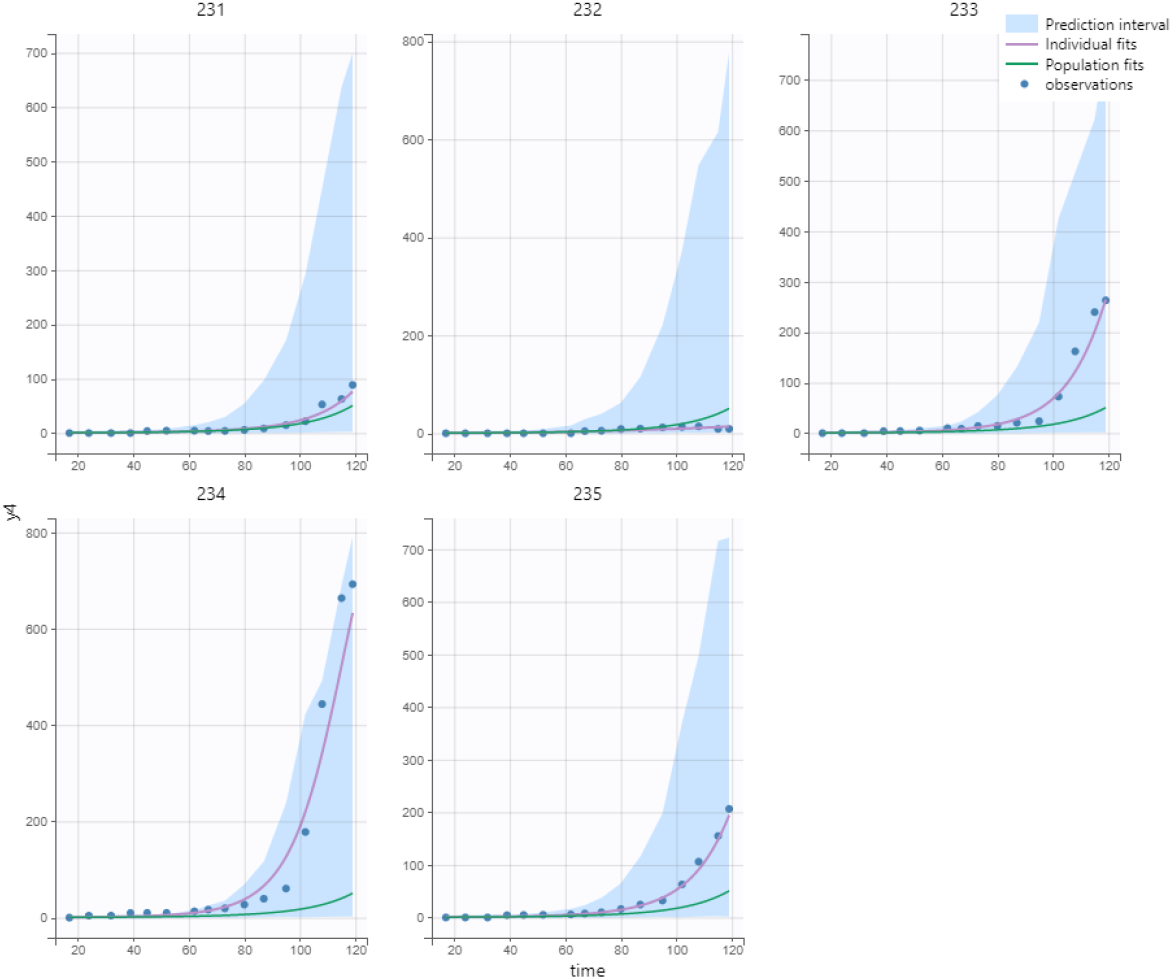
Simulation results for *T*(*t*) in case of low E2 level in Model (5).

**Figure 23:**
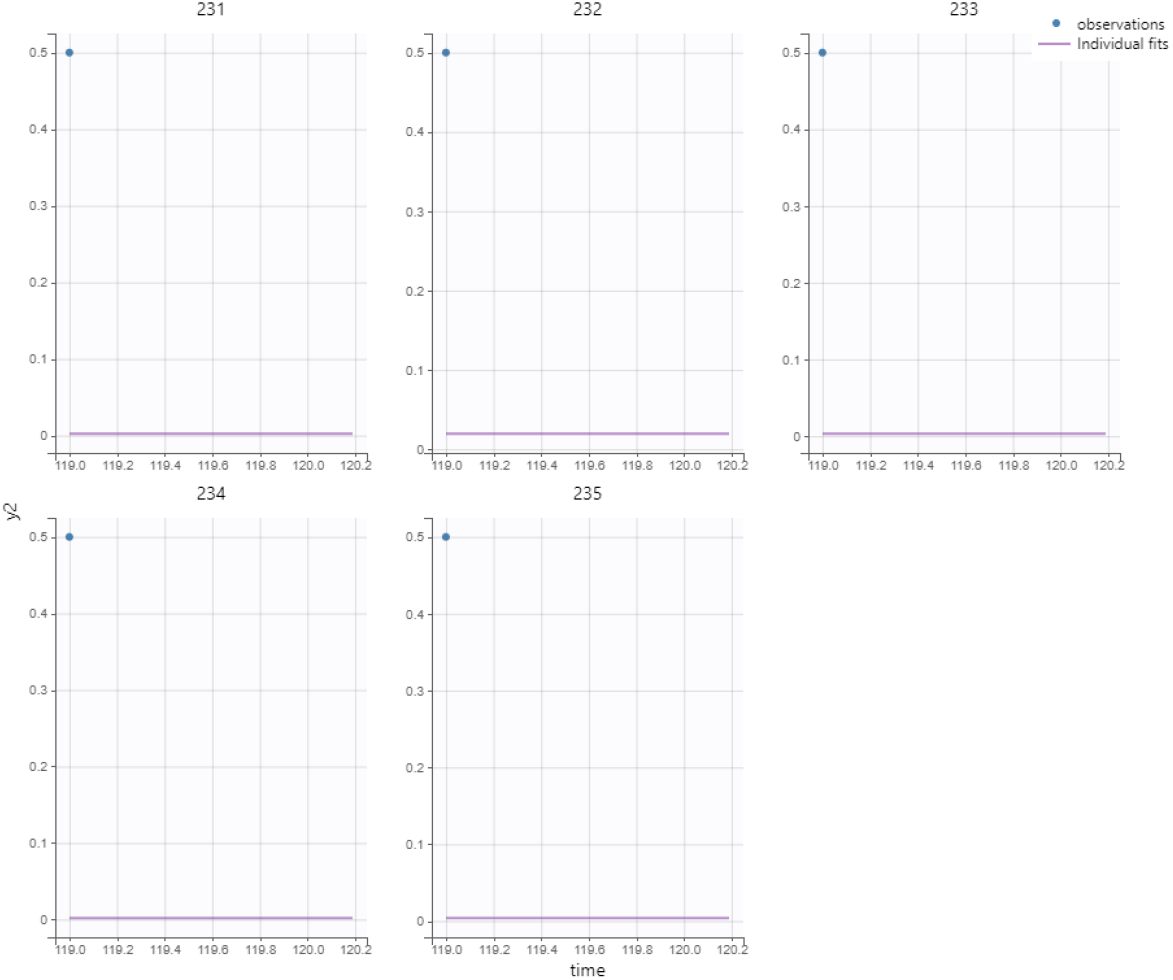
Model calibration for *ER*(*t*) in case of high E2 level in Model (5).

### Plots for observations vs predictions

**Figure 24:**
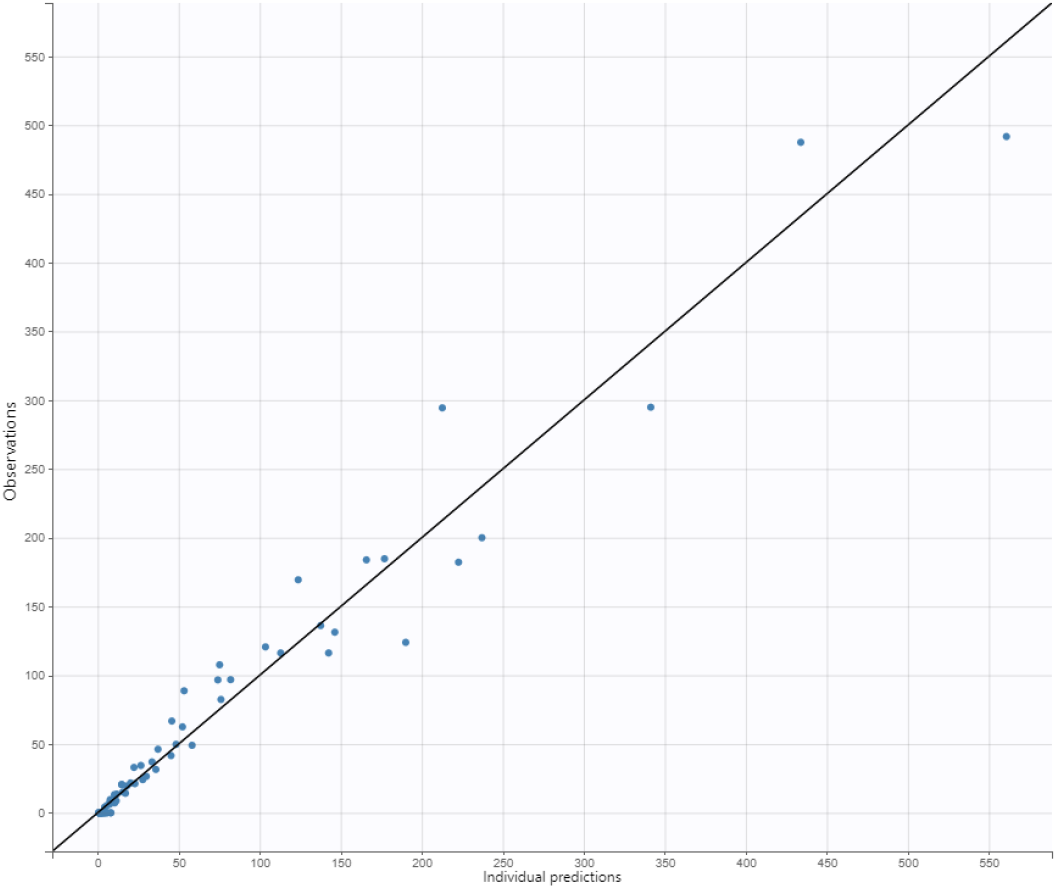
Observations vs predictions for *T*(*t*) in case of high E2 level in Model (1).

**Figure 25:**
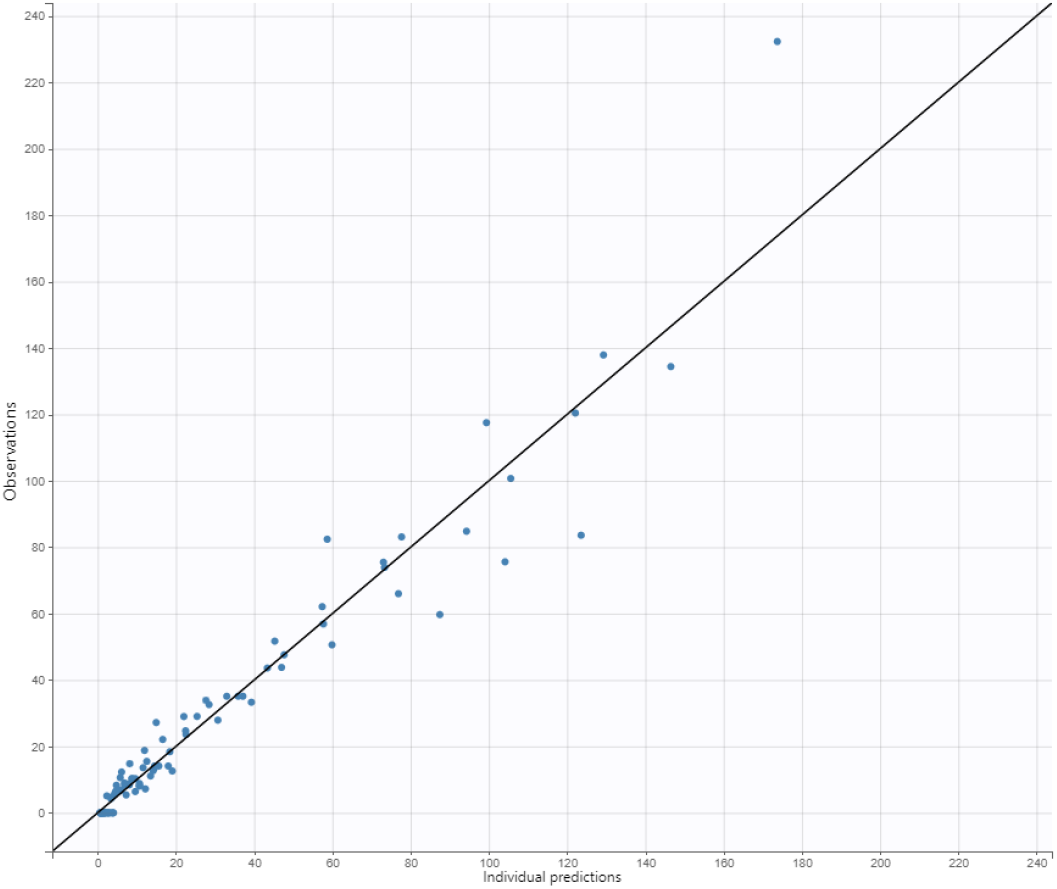
Observations vs predictions for *T*(*t*) in case of medium E2 level in Model (1).

**Figure 26:**
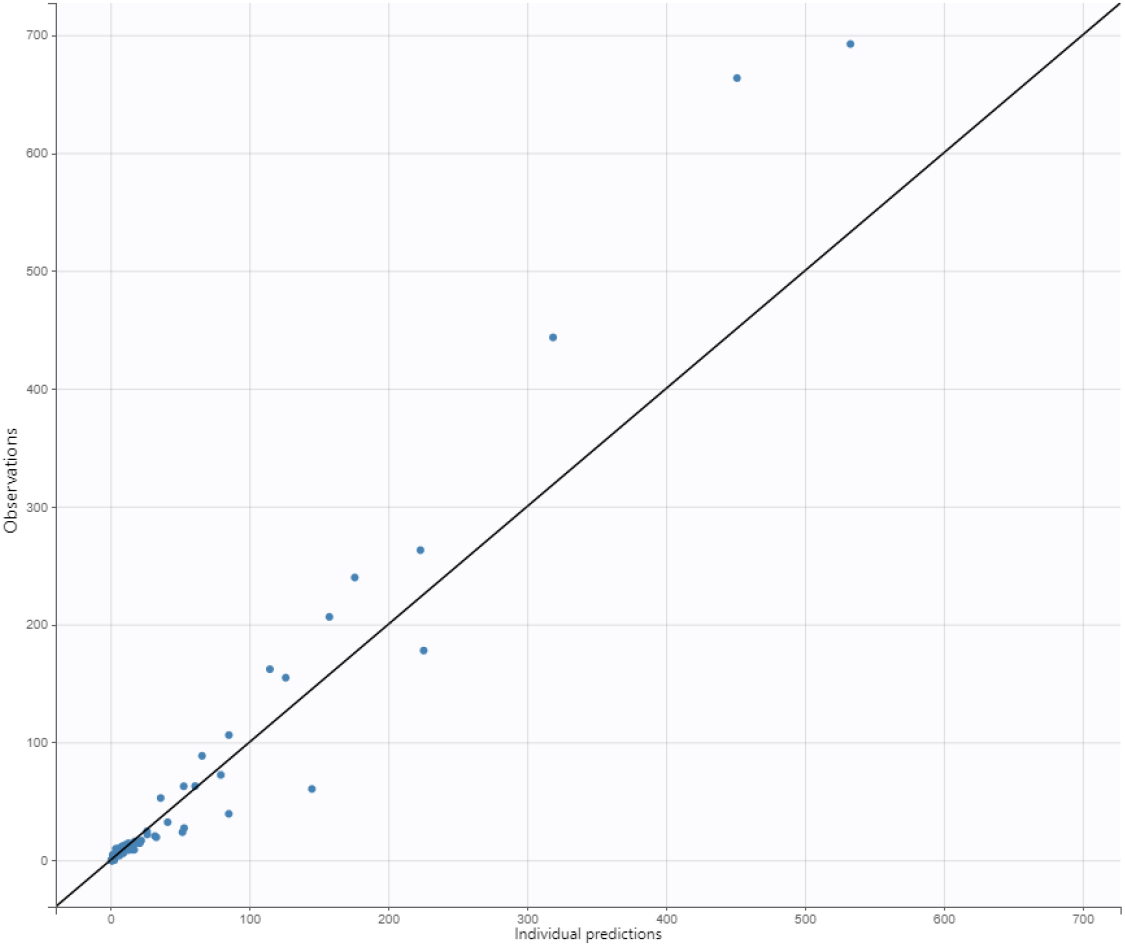
Observations vs predictions for *T*(*t*) in case of low E2 level in Model (1).

**Figure 27:**
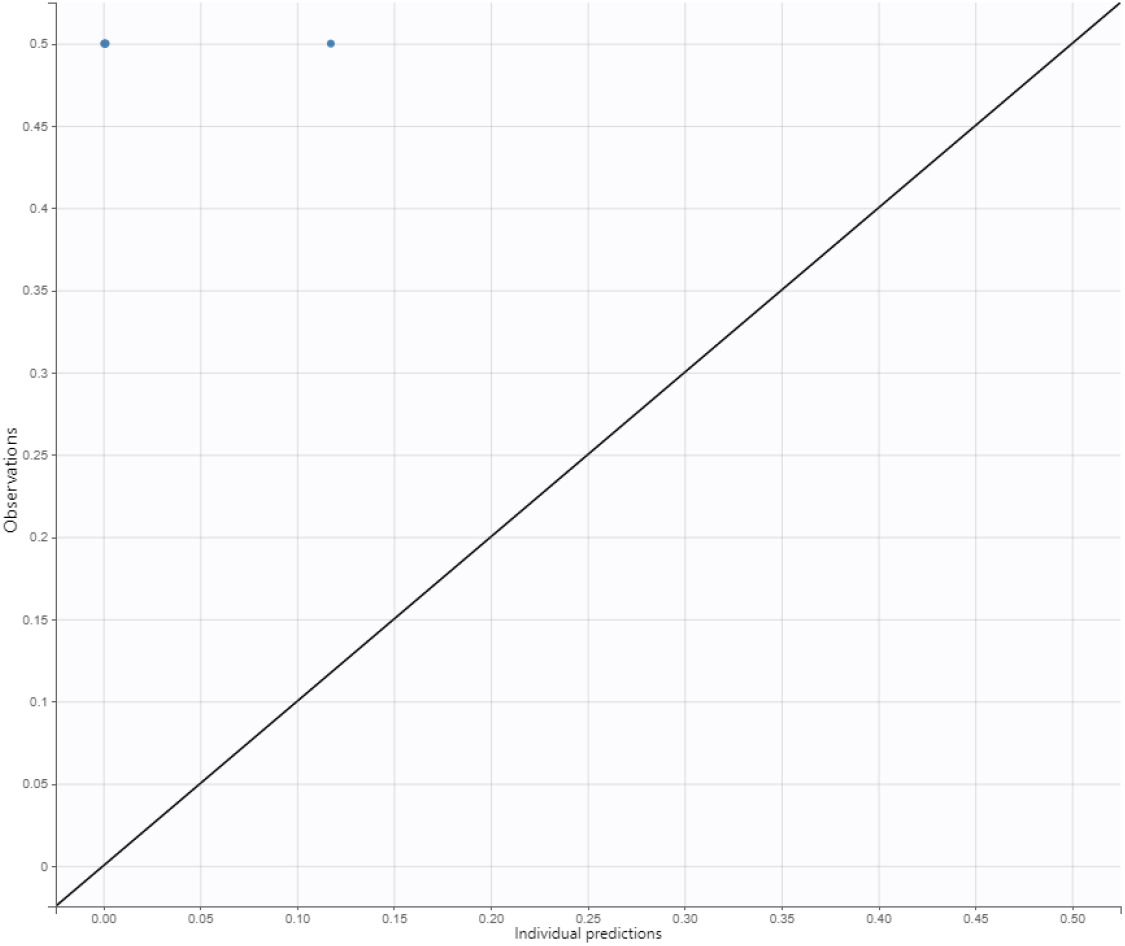
Observations vs predictions for *ER*(*t*) in case of high E2 level in Model (1).

**Figure 28:**
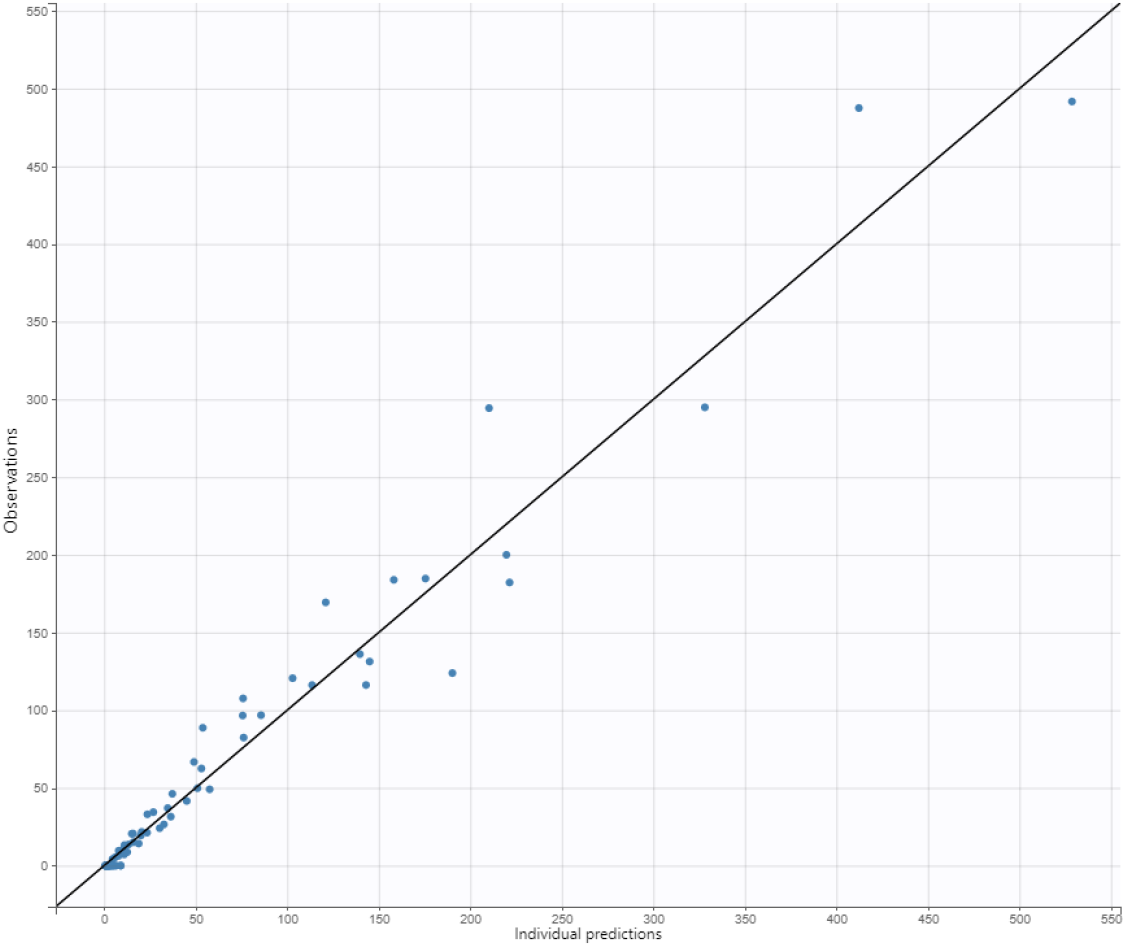
Observations vs predictions for *T*(*t*) in case of high E2 level in Model (5).

**Figure 29:**
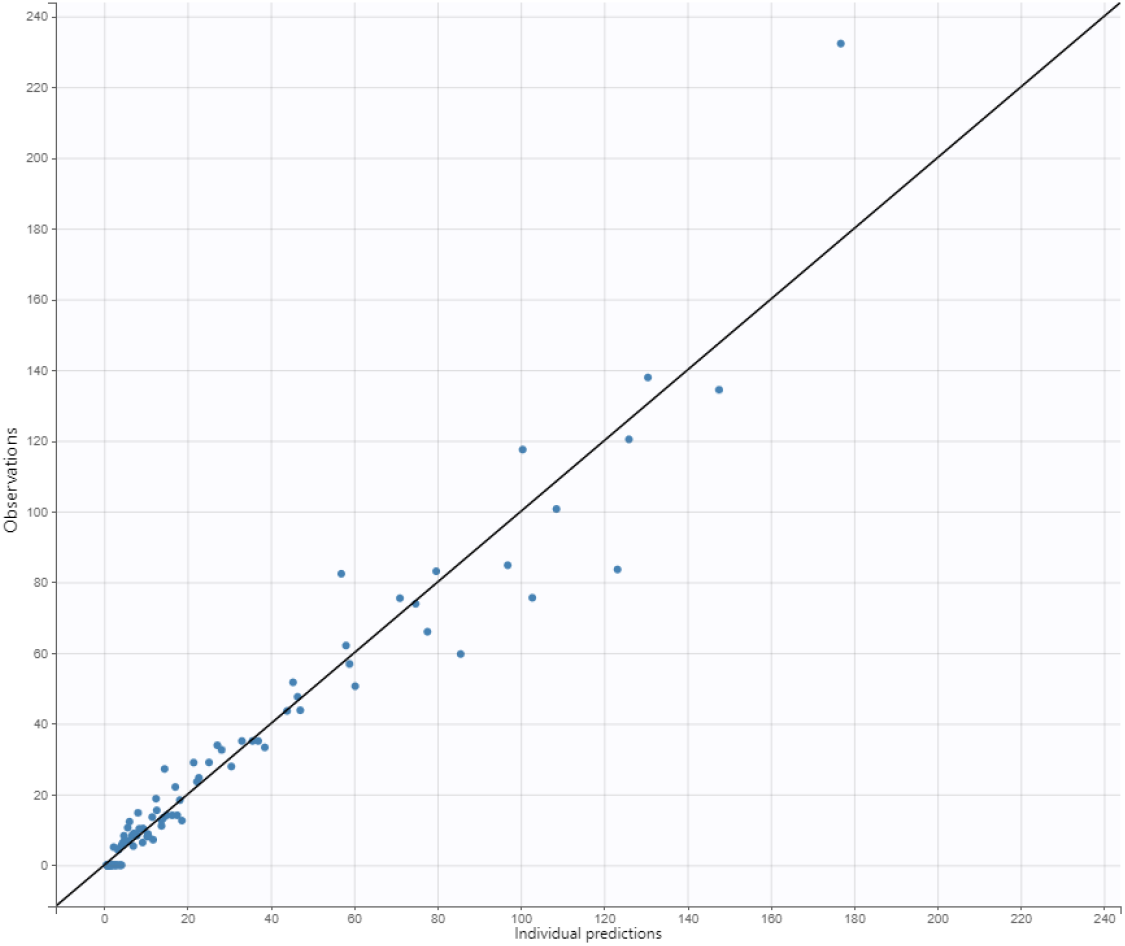
Observations vs predictions for *T*(*t*) in case of medium E2 level in Model (5).

**Figure 30:**
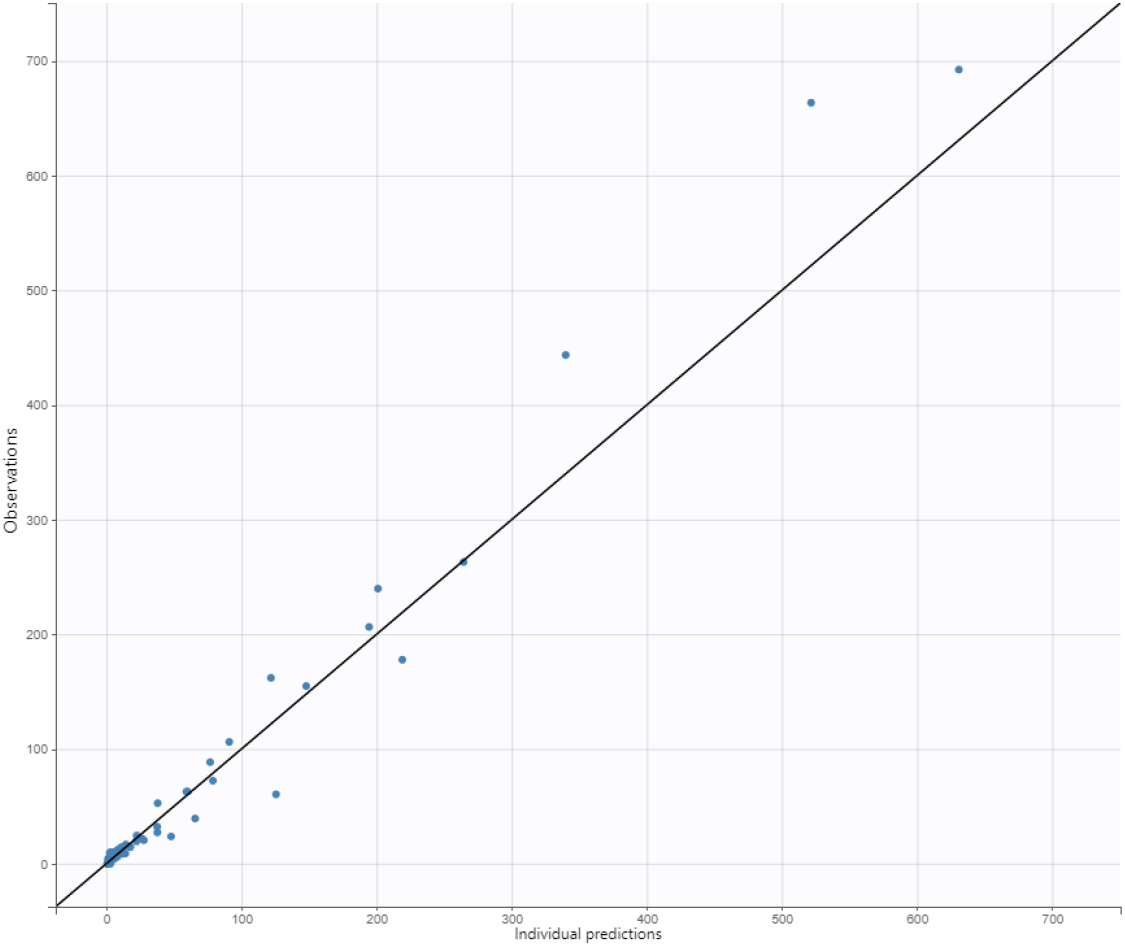
Observations vs predictions for *T*(*t*) in case of low E2 level in Model (5).

**Figure 31:**
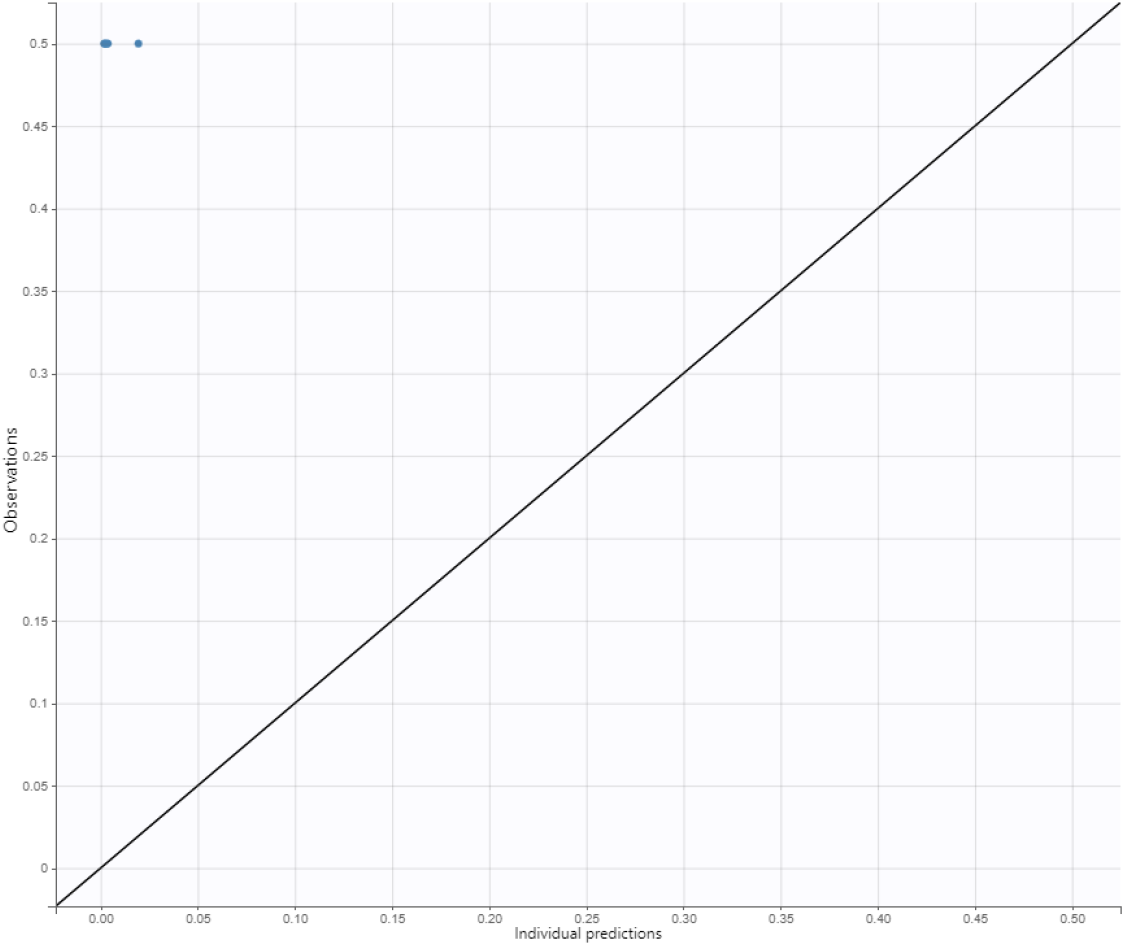
Observations vs predictions for *ER*(*t*) in case of high E2 level in Model (5).

## Ethics

All animal studies were approved by the local ethics committee for animal experimentation in Lund / Malmö, under permit# 13939-25 and 14122-2020.

## Data Access

Data and code are available in https://github.com/TugbaAkmanTR/CAF-TARGET.

## Author contribution

TA: conceptualization, formal analysis, investigation, methodology, project administration, supervision, validation, software, visualization, writing—original draft, writing—review and editing; KP: Data curation, resources, Writing – review and editing; AA: conceptualization, investigation, project administration, supervision, writing—original draft, writing—review and editing; AKL: investigation, methodology, supervision, writing—review and editing.

All authors gave final approval for publication and agreed to be held accountable for the work performed therein.

## Acknowledgmen

The authors acknowledge the use of artificial intelligence–based tools to support code development and to refine the presentation of tables and figures. The AI tools did not contribute to the scientific content or interpretation of the results.

## Disclaimer

The authors have no disclaimers to declare.

